# A buffer-tuning strategy to profile domain-specific activity of chimeric I-TevI/CRISPR gene editors *in vitro*

**DOI:** 10.1101/2025.10.24.684378

**Authors:** Kurt W. Loedige, Alexa L. White, Thomas A. McMurrough, Brent E. Stead, David R. Edgell

## Abstract

Protein-DNA interactions can be manipulated *in vitro* by changing buffer conditions. Here, we develop a methodology to map the cleavage preferences of chimeric gene editors that are fusions of the I-TevI nuclease domain to CRIPSR nucleases by manipulating *in vitro* salt concentrations. We found that DNA cleavage by the I-TevI (Tev) nuclease domain at CNNNG sites was de-coupled from the gRNA-targeted site in low salt buffers. For TevCas12a, this non-targeted cleavage activity was enriched at Tev CNNNG cleavage motifs optimally positioned within a 30-bp window upstream of a Cas12a TTTV PAM site. Non-targeted cleavage did not require Cas12a nuclease activity or specific Cas12a gRNA targeting. Similar non-targeted products were observed in low salt buffer conditions for TevSaCas9, Tev-meganuclease and Tev-zinc finger editors. Cas12a and SaCas9 activity at gRNA-directed sites and sites with multiple mismatches were also sensitive to buffer salt concentration. Oxford Nanopore sequencing revealed a remarkably similar Tev CNNNG cleavage preference at different salt concentrations and in different fusion contexts, emphasizing the robustness and specificity of Tev activity. More generally, our work highlights the sensitivity of gene editors to *in vitro* reaction conditions and how these conditions can be leveraged to functionally dissect the activity of individual domains of chimeric gene editors.

## Introduction

Genome editing nucleases are programmable, site-specific DNA endonucleases that enable targeted genome modification^1^, ^2^. A critical aspect of their application is understanding DNA binding and cleavage specificity, which informs the selection of on-target sites that support high activity while minimizing activity at mismatch-containing off-target sites. Profiling the specificity of gene editors often relies on *in vitro* methodologies that allow activity determination on a larger number substrates than is possible *in vivo*^3–5^. However, *in vitro* protein-DNA interactions are sensitive to buffer conditions^6–9^. In particular, modulating ionic buffer strength has been leveraged as a tool to probe the mechanisms and kinetics of gene editors, including zinc-finger nucleases^10^, TALE domains^11^, ^12^, Cas9^13^, ^14^, and Cas12a^15^.

We previously showed that the GIY-YIG nuclease domain and interdomain linker from the homing endonuclease I-TevI (Tev) could be fused to unrelated DNA-binding modules (zinc fingers, TALEs, meganucleases and Cas9 orthologs) to create chimeric gene editors^16–20^. In these editors, the DNA-binding module dictates target site selection, while Tev cleaves at a CNNNG motif that is appropriately spaced from the target site, with some differences in spacing for each fusion context^16–18^. In particular, the fusion of Tev to SpCas9 from *Streptococcus pyogenes*, a class II type II-B nuclease^19^, created an

RNA-guided dual nuclease (TevSpCas9) that makes two double-strand breaks (DSBs): one at a Tev CNNNG motif upstream of the 5’ end of the gRNA target site (producing a 2-nt 3’ overhang) and another at the Cas9 cut site (yielding a blunt end). In mammalian cells, TevSpCas9 editing generates defined length deletions with high frequency and no detectable exonucleolytic processing of the DNA ends^19^.

We sought to expand on the dual RNA-guided nuclease architecture by fusing Tev to the N-terminus of *Acidaminococcus sp*. Cas12a (TevCas12a) (Fig. 1A). Cas12a (Cpf1) is a class II, type V-A CRISPR nuclease that cleaves double-strand DNA (dsDNA) at sites specified by a 20-nucleotide crRNA adjacent to a TTTV (V=A/C/G) protospacer-associated motif (PAM) sequence^21^, ^22^. Cas12a and related proteins have been adapted for gene editing in mammalian systems and for nucleic acid diagnostic applications^23–26^. We anticipated that cleavage by TevCas12a would generate two DSBs: one by Tev cleavage at a CNNNG motif to create a 2-nt 3’ overhang, and the other by Cas12a to create a 4-nt 5’ overhang.

**Figure 1.**
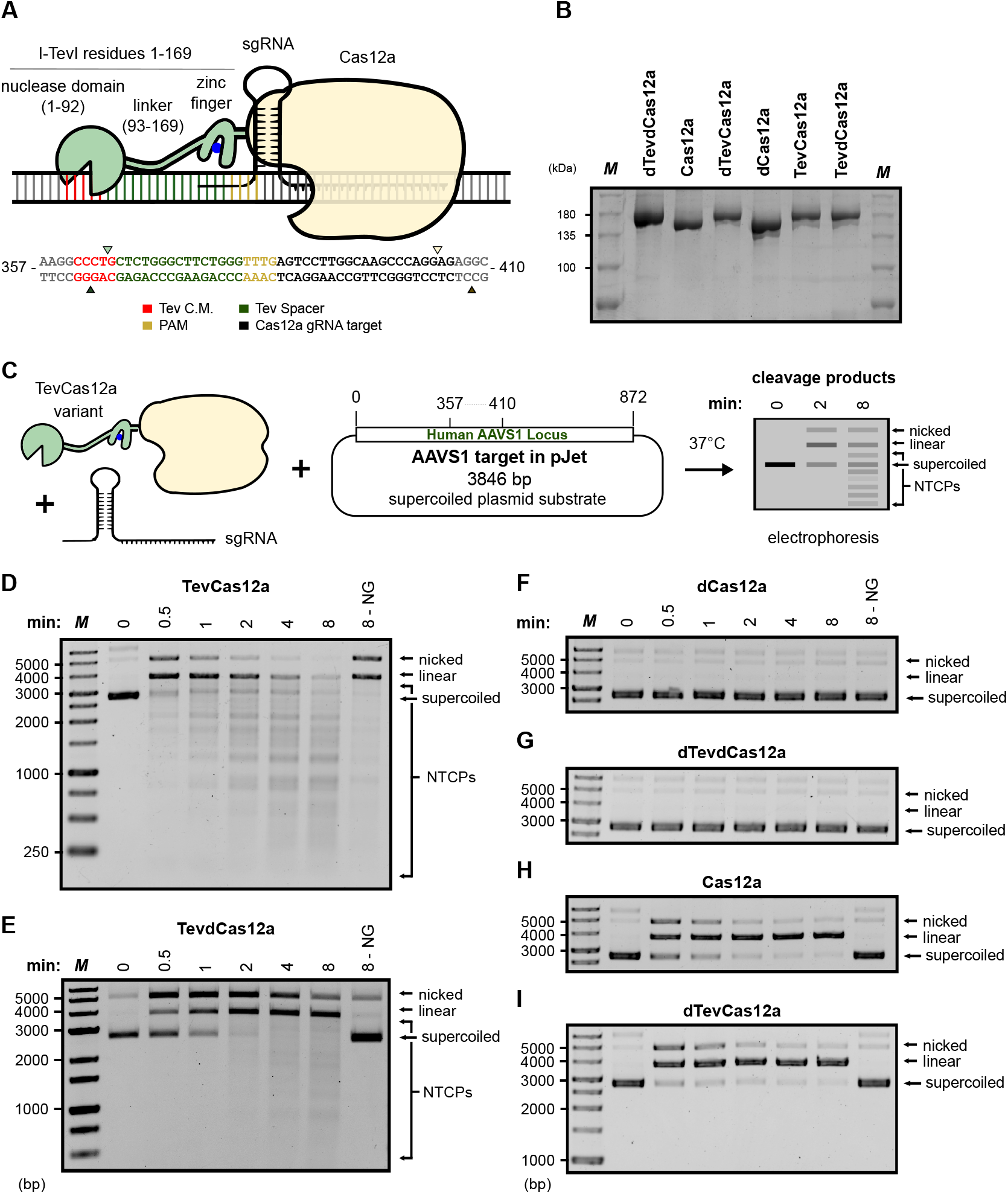
Non-targeted cleavage of supercoiled DNA by TevCas12a. **(A)** Schematic diagram of TevCas12a with annotated components alongside the pJET-AAVS1 target site sequence. Triangles indicated Tev or Cas12a cleavage sites. **(B)** Purified TevCas12a and Cas12a protein variants. “d” represents catalytically inactive (dead) domains. *M*, molecular marker with sizes indicated on the left of the gel. **(C)** Schematic diagram of *in vitro* cleavage reactions: purified proteins are formed into RNPs by incubation with targeted or non-targeted gRNA, incubated with supercoiled plasmid at 37°C and products resolved by agarose gel electrophoresis. **(D)** Time-course cleavage of pJET-AAVS1 with TevCas12a/gRNA_AAVS1_ showing both targeted Cas12a cleavage products and non-targeted cleavage products (NTCPs). M, molecular weight ladder with sizes in bp indicated on left. NG, no guide. **(E-I)** Time-course cleavage assays with pJET-AAVS1 and (E) TevdCas12a, (F) dCas12a, (G) dTevdCas12a, (H) Cas12a, or (I) dTevCas12a complexed with gRNA_AAVS1_. Labeled as in panel D. Equivalent time-course assays with a non-targeted gRNA (gRNA_CFTR_) are provided in Supplementary Fig. S2A–B. Uncropped gel images for panels B-I are shown in Supplementary Fig. S7.

Here, we characterize TevCas12a activity *in vitro* and unexpectedly find that dsDNA cleavage activity is uncoupled from the gRNA-targeted site in specific, low salt buffer conditions. This non-targeted cleavage activity requires Tev activity and Cas12a DNA binding, but not Cas12a nuclease activity. Using target site knockouts and Oxford Nanopore sequencing, we show that TevCas12a cleaves substrates at Tev CNNNG motifs that are optimally spaced 20-bp from a Cas12a TTTV PAM site. Non-targeted cleavage products were also observed *in vitro* with TevSaCas9, Tev-zinc finger nucleases, and Tev-meganuclease fusions. Our results demonstrate how Tev and Cas12a use different strategies to search for target sites and how these activities can be leveraged to map cleavage preferences of chimeric gene editors.

## Results

### Non-targeted cleavage of dsDNA substrates by TevCas12a

To construct TevCas12a, we fused I-TevI residues 1-169 encompassing the GIY-YIG nuclease domain and interdomain linker to the N-terminus of *Acidaminococcus sp*. Cas12a (TevCas12a), as well as to its catalytically dead variant D908P (TevdCas12a) (Fig. 1A,B). During our initial characterization of TevCas12a, we noted that the cleavage activity was sensitive to *in vitro* buffer conditions. In particular, low cleavage activity was observed in reactions with greater than 100 mM sodium chloride. To further characterize TevCas12a activity, we created a supercoiled plasmid substrate (pJET-AAVS1) with a ∼1kb insert of the AAVS1 safe harbour site that was PCR amplified from human genomic DNA (Fig. 1C). When pJET-AAVS1 was incubated with RNPs consisting of purified TevCas12a (Fig. 1B) and an *in vitro* transcribed gRNA_AAVS1_ in buffer containing 50 mM sodium chloride and 1 × rCutSmart buffer, we observed robust on-target activity as well as multiple cleavage products ranging in size from ∼100-3000-bp (Fig. 1D). These products were inconsistent with the size of the predicted TevCas12a linear products based on the position of the gRNA_AAVS1_ target site in the *AAVS1* sequence (3.801-kb and a small 0.045-kb product corresponding to the fragment between the Tev and Cas12a cleavage sites) (Fig. 1C). We termed these cleavage products non-targeted cleavage products (NTCPs). At the end of the 8 min timecourse at a TevCas12a:pJET-AAVS1 ratio of 20:1, *>*95% of pJET-AAVS1 was converted to NTCPs (Fig. 1D). Lowering the TevCas12a:pJET-AAVS1 ratio reduced the appearance of NTCPs (Fig. S1E-J).

Interestingly, we also observed cleavage of pJET-AAVS1 into a discrete linear product and NTCPs with TevCas12a in the absence of the gRNA_AAVS1_ (apo-TevCas12a) after 8 min, albeit at a lower rate than when reactions were performed with TevCas12a/gRNA_AAVS1_ (Fig. 1D, NG (no guide)). NTCPs were also observed when we targeted TevCas12a to isolated linear *λ* DNA with a gRNA targeting gene *N* (gRNA_*λ*_) (Fig. 2A) or to a lesser extent with apo-TevCas12a at high protein:DNA ratios (Fig. 2C). The non-targeted activity of TevCas12a was enhanced in reactions containing 1 *×* CutSmart where sodium chloride concentrations were 100 mM or less, whereas the on-target gRNA-directed cleavage fraction was highest in buffers with up 200 mM sodium chloride concentrations (Fig. 2A and B). As rCutSmart (NEBuffer 4) contains 50 mM potassium acetate (K^+^), these sodium chloride concentrations represent additive increases in total monovalent cations (e.g., 100 mM Na^+^ corresponds to 150 mM total monovalent cations).

**Figure 2.**
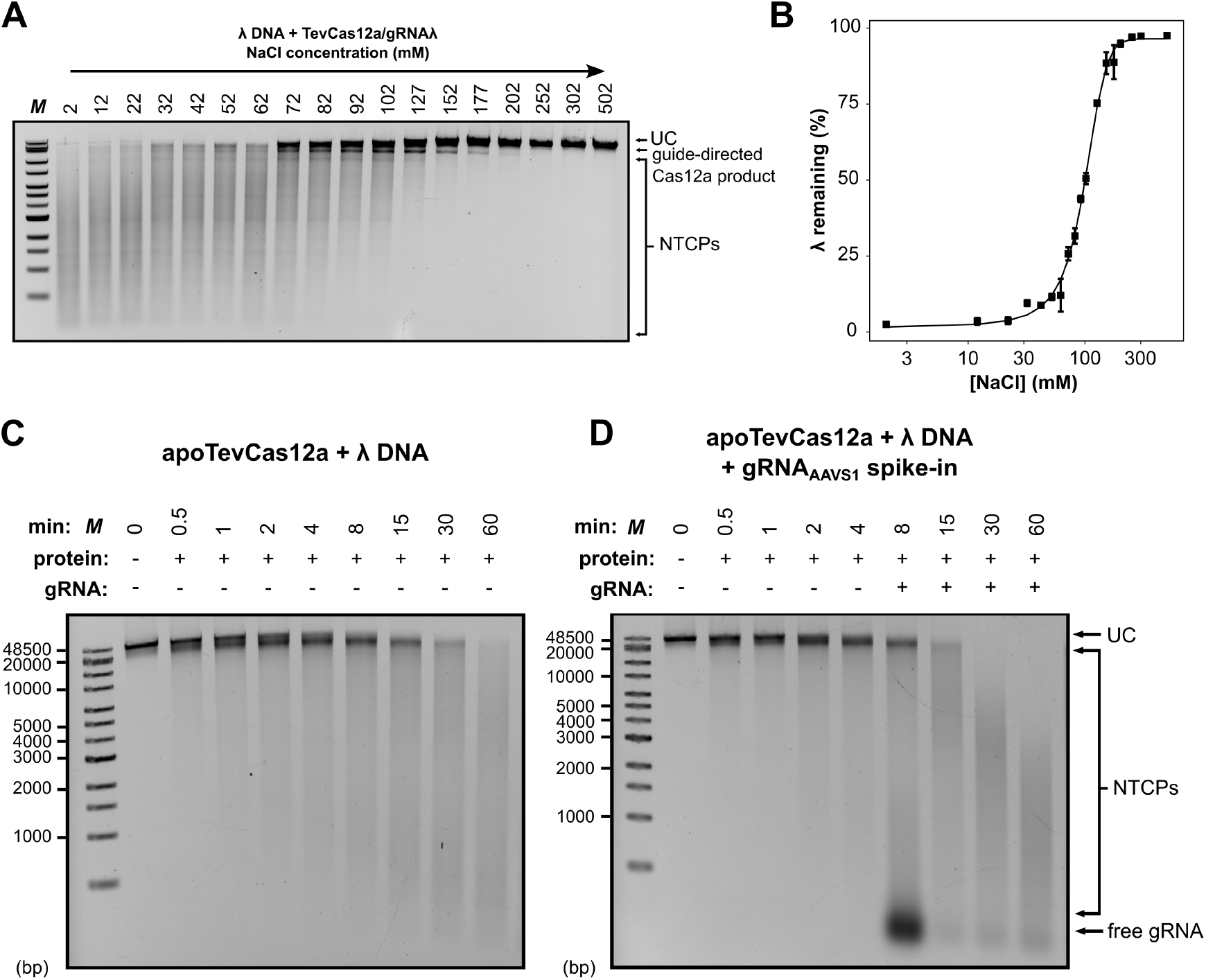
Effect of NaCl concentration on CutSmart-buffered TevCas12a cleavage of *λ* DNA. **(A)** Representative agarose gel of titration of sodium chloride concentration in reactions with *λ* DNA and TevCas12a/gRNA_*λ*_. The final sodium chloride concentration in each reaction is indicated above the indicated lane. Products and unreacted *λ* genome (UC) are indicated on the right of the gel. M, molecular weight marker. NTCPs, non-targeted cleavage products. **(B)** Plot of reaction sodium chloride concentration versus unreacted *λ* DNA remaining. Each point is mean of three independent replicates with error bars indicating standard deviation from the mean. The data were fit to a non-linear regression model. **(C)** Reaction of apoTevCas12a and *λ* DNA at the indicated time points. **(D)** Spike-in of gRNA to reaction of apoTevCas12a and *λ* DNA. Uncropped gel images for panels A-D are shown in Supplementary Fig. S8.

Cleavage of dsDNA by Cas12a proceeds by nicking of the non-targeted DNA strand followed by conformational rearrangement of the RuvC domain to nick the targeted DNA strand^21^, ^22^. With a super-coiled plasmid substrate, these steps are visible as a nicked intermediate and linear product, and can be estimated by first-order reaction rates *k*_1_ and *k*_2_, respectively (Table 1, Fig. S1A–D). However, with an active TevCas12a fusion that generates multiple NTCPs, individual nicking reaction products are difficult to distinguish and thus the overall reaction rate (*k*_*obs*_) is estimated from depletion of supercoiled plasmid. Notably, TevCas12a timecourses were performed at a 10:1 enzyme:substrate ratio, whereas Cas12a, TevdCas12a and dTevCas12a timecourses were performed at 20:1, so the reported *k*_*obs*_ likely underestimates the true TevCas12a reaction rate. Even under these conditions, the TevCas12a *k*_*obs*_ rate (0.112 *±* 0.010 nM min^−1^) is very similar to the Cas12a *k*_1_ rate (0.118 *±* 0.016 nM min^−1^) (Table 1). To help distinguish NTCPs from Cas12a on-target cleavage, we used the TevdCas12a fusion (Figs. 1E, S1B) and found that the *k*_*obs*_ rate was reduced by ∼1.8-fold (0.067 *±* 0.009 nM min^−1^). The difference in reaction rates for TevdCas12a versus TevCas12a could be related to the extremely slow dissociation rate of dCas12a from dsDNA in Mg^2+^-containing buffer^27^. Profiling the temperature dependence revealed a reduction of 50% and 75% in the appearance of non-targeted products at 30°C and 18°C as compared to 37°C (Fig. S2C).

**Table 1.** Observed reaction rates for targeted (*k*_*1*_ and *k*_*2*_) and non-targeted (*k*_*obs*_) cleavage of pJET-AAVS1 (nM DNA/sec) for all proteins at 20:1, except TevCas12a at 10:1 (*) protein:DNA.

| Enzyme | $k_1$ | $k_2$ | $k_{obs}$ |
| --- | --- | --- | --- |
| Cas12a | $0.118 \pm 0.016$ | $0.035 \pm 0.007$ | N.D. |
| TevCas12a* | N.D. | N.D. | $0.112 \pm 0.010$ |
| TevdCas12a | N.D. | N.D. | $0.067 \pm 0.009$ |
| dTevCas12a | $0.053 \pm 0.011$ | $0.038 \pm 0.011$ | N.D. |

### Non-targeted cleavage is indepdendent of gRNA targeting

To test the dependence of NTCPs on specific gRNA-target DNA interactions, we designed a gRNA to target a site in the human cystic fibrosis transmembrane conductance regulator (*CFTR*). When TevCas12a/gRNA_CFTR_ or TevdCas12a/gRNA_CFTR_ was incubated with pJET-AAVS1, we observed NTCPs at approximately the same rate as with the gRNA_AAVS1_ (Fig. S2A-B, compare with Fig. 1D-E). Spiking gRNA_AAVS1_ into a reaction of apo-TevCas12a and *λ* DNA at 8 min stimulated the appearance of NTCPs (Fig. 2D, compare with Fig. 2C). This result shows that gRNA stimulates NTCP generation, but that the stimulatory effect is not dependent on targeted gRNA-DNA interactions. To further illustrate this effect, we quantified *λ* DNA conversion to NTCPs across the three conditions: apo-TevCas12a (Fig.2C), apo-TevCas12a with gRNA_AAVS1_ spiked in at 8 min (Fig. 2D), and TevCas12a with a non-targeted gRNA_AAVS1_ included from the start (Fig. 4D). The resulting data (Fig. S3) confirm that both spiked and non-targeted gRNAs reproducibly stimulate NTCP formation relative to apo-TevCas12a. The decrease of free gRNA at later time points reflects the incorporation of gRNA into RNP complexes over the course of the reaction. Moreover, the appearance of multiple NTCPs cannot be explained by cleavage of TevCas12a/gRNA_AAVS1/CFTR_ at off-target sites. Previous work has shown that Cas12a can nick dsDNA with gRNA-target mismatches of 4 or less^28^; there are no sites with 6 or fewer mismatches in pJET-AAVS1 to gRNA_CFTR_, and no sites with 7 or fewer mismatches to gRNA_AAVS1_.

### Non-targeted cleavage requires Tev but not Cas12a nuclease activity

We next generated an active site mutant in the Tev nuclease domain (R27A, dTev)^29^, ^30^ to determine the dependency of NTCPs on Tev nuclease activity. With a TevdCas12a mutant (Fig. 1B, E), we found a reduction in the rate and extent of NTCPs (Fig. S1B). Nicked and linear products consistent with gRNA-targeted cleavage at the AAVS1 site by the Tev domain were still observed (Fig. 1E). No cleavage products were observed with a dCas12a mutant (Fig. 1B, F). However, with the reciprocal dTevCas12a mutant (Fig. 1B, I) that knocked out the Tev active site, NTCPs were abolished while on-target gRNA directed Cas12a cleavage was not impacted (Fig. 1I, Fig. S2D). The double mutant, dTevdCas12a (Fig. 1B), abolished both NTCPs and on-target gRNA cleavage (Fig. 1G). The appearance of NTCPs with the non-targeted *CFTR* gRNA was also dependent on an active Tev nuclease domain and were reduced with the TevdCas12a mutant, similar to that observed with gRNA_AAVS1_ (Fig. S2A-B).

Taken together, these data show that by simply changing *in vitro* salt concentration, TevCas12a activity can be modified to a non-targeted cleavage activity that does not require a programmed gRNA-DNA interaction. The non-targeted products and specific Cas12a-gRNA directed products appear at high protein:DNA ratios, consistent with single-turnover kinetics. The differences in salt sensitivity for the NTCPs and the gRNA-directed products suggest that these two activities result from different DNA cleavage events. Inactivation of Tev nuclease activity abolishes the appearance of NTCPs, whereas inactivation of Cas12a nuclease reduces but does not abolish the appearance of NTCPs. These observations rule out Cas12a collateral cleavage activity as a source of NTCPs. Moreover, the appearance of discrete products rather than complete plasmid degradation suggests that the non-targeted cleavage activity exhibits sequence specificity, consistent with the CNNNG preference of the Tev nuclease domain.

### Tev CNNNG motifs are required for non-targeted cleavage

To further dissect the DNA target sequence requirements for the NTCPs, we next created a series of 500-bp substrates derived from pJET-AAVS1 where we mutated the Tev CNNNG cleavage motifs (ΔCNNNG) to non-cleavable sites by mutating the first C of the motif, the Cas12a TTTV PAM sites (ΔTTTV) by mutating the first T of the PAM, or both (Fig. 3A). In all cases, the on-target gRNA binding site, TTTV PAM and Tev CNNNG site (except for ΔCNNNG substrate) were retained. As shown in Fig. 3B with the wild-type substrate, the predicted on-target gRNA directed cleavage products and NTCPs were observed at 0.5 min, with complete substrate cleavage by 15 min. With the ΔCNNNG substrate that knocked out all CNNNGs, NTCPs were not observed (Fig. 3C). Interestingly, NTCPs were observed with the ΔTTTV substrate (Fig. 3D), although to a lesser extent and differently patterned than with the wild-type substrate (Fig. 3B). NTCPs and the Tev cleavage product 1 (TP1) were observed in the absence of an added gRNA, indicating that apo-TevCas12a retains DNA-binding activity at the protein:DNA concentrations used in the experiment (Fig. 3B, D). With the double ΔCNNNG + ΔTTTV knockouts, we observed only the on-target Tev and Cas12a cleavage products (Fig. 3E). Collectively, these data show that NTCPs require Tev CNNNG motifs and are enhanced with substrates that also contain Cas12a TTTV PAM sites.

**Figure 3.**
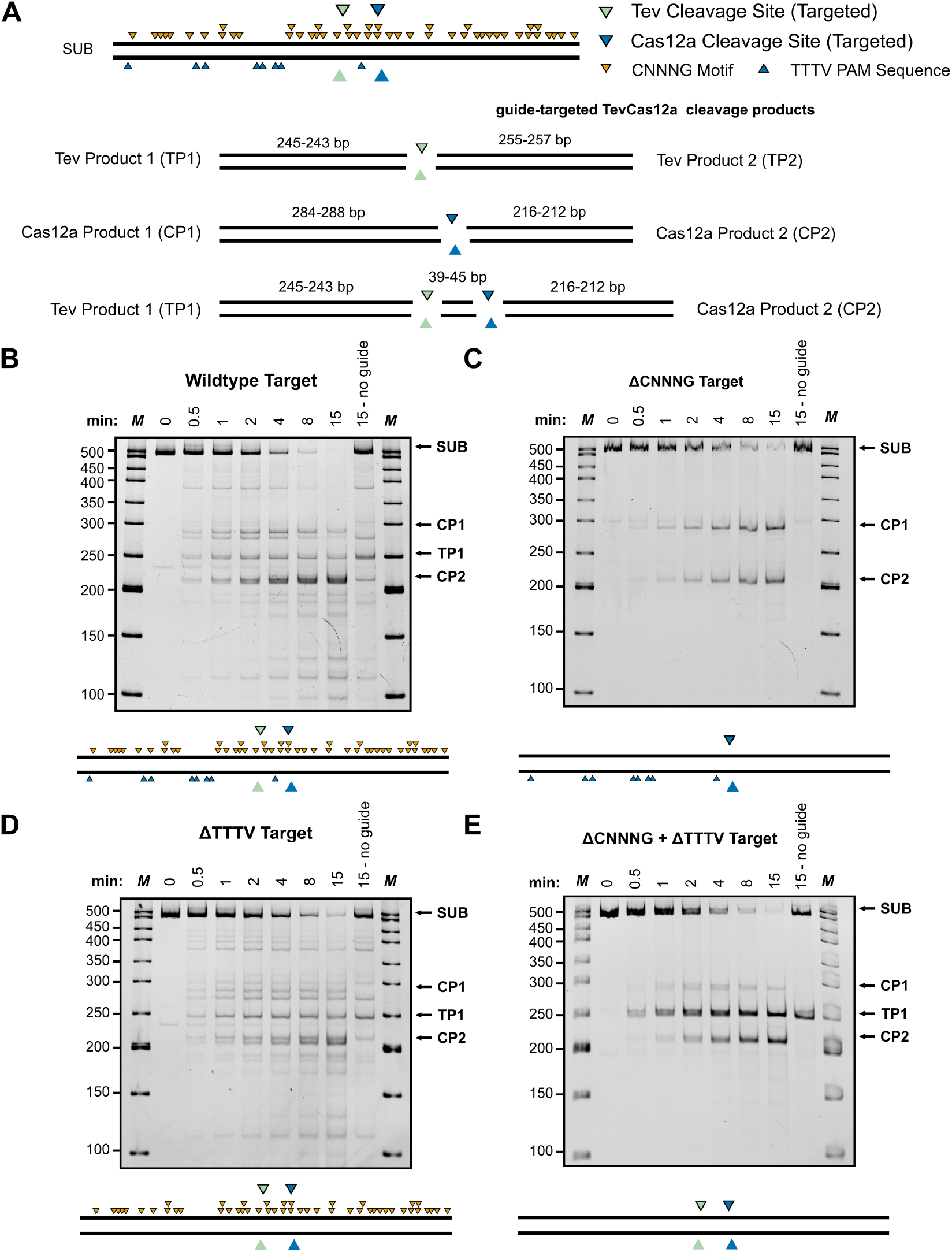
TevCas12a cleavage of wildtype and mutated AAVS1 substrates. **(A)** Schematic of the linear wild-type AAVS1 substrate (SUB) indicating gRNA targeted Tev and Cas12a cleavage sites and non-targeted Tev CNNNG cleavage motifs and Cas12a TTTV PAM sequences. For simplicity, CNNNG motifs are depicted on one strand and TTTV PAM sites on the other strand. Predicted guide-targeted cleavage products are shown below. **(B-E)** TevCas12a cleavage timecourse assays on wild-type, ΔCNNNG, ΔTTTV and ΔCNNNG/ΔTTTV substrates. For each gel image, unreacted substrate (SUB) and targeted Tev (TP1) and Cas12a (CP1-2) cleavage products are annotated. Uncropped gel images for panels B-E are shown in Supplementary Fig. S9.

**Figure 4.**
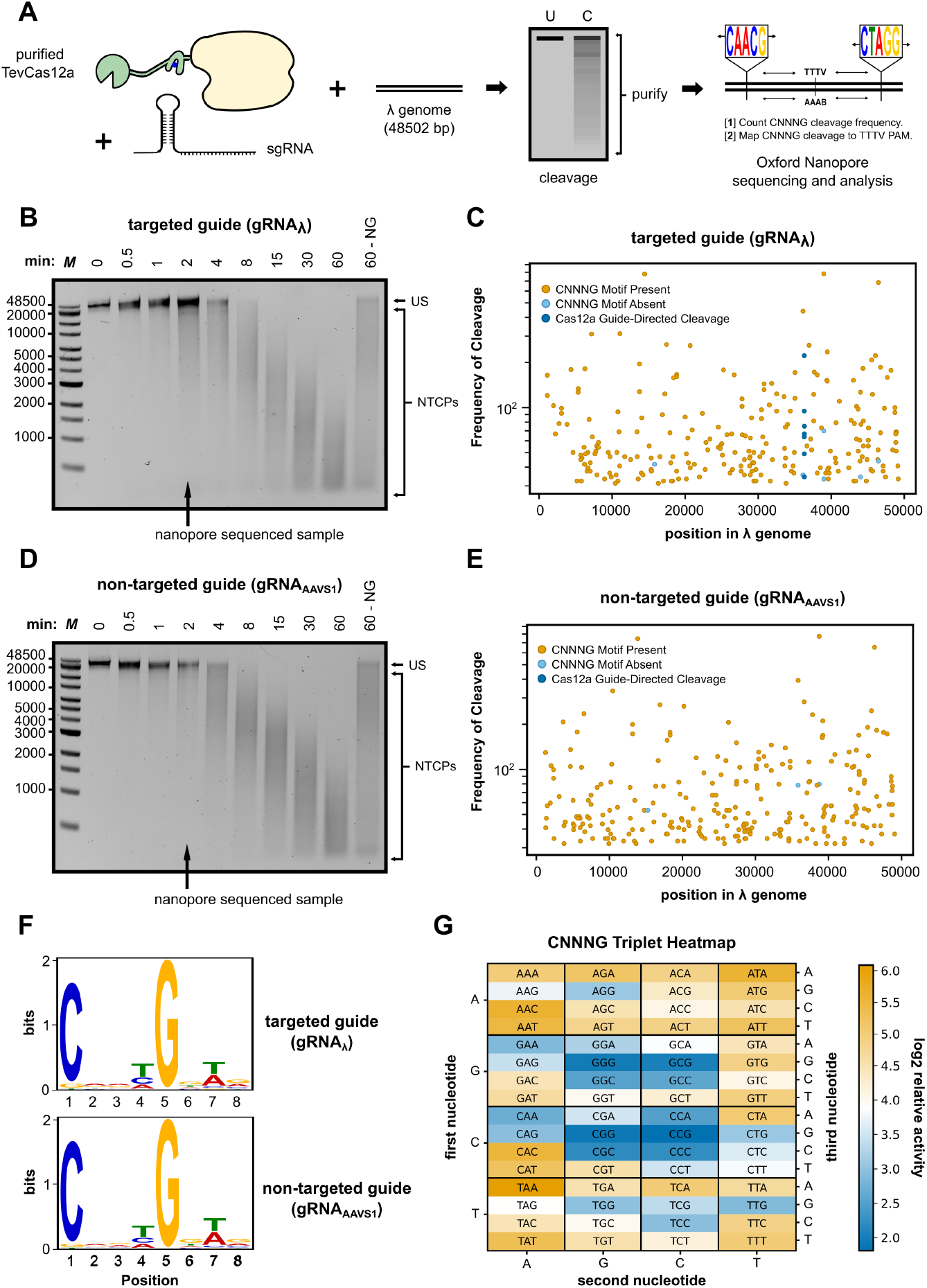
Mapping TevCas12a non-targeted cleavage using Oxford Nanopore sequencing. **(A)** Schematic overview of the experimental workflow. Purified TevCas12a and guide RNA (sgRNA) were incubated with linear *λ* DNA, followed by cleavage and purification of NTCPs. Sequenced DNA fragments were mapped by position, surrounding sequence was extracted, and cleavage events were counted and measured relative to TTTV PAMs. **(B**,**D)** Agarose gel of TevCas12a *λ* genome cleavage time course with a targeted (gRNA_*λ*_ or non-targeted (gRNA_AAVS1_) guide. US: uncleaved substrate; NTCPs: non-target cleavage products. **(C**,**E)** Scatterplot depicting the frequency and distribution of top 250 cleavage events across the *λ* genome. Each dot represents a cleavage position, colored by CNNNG motif presence or predicted Cas12a-directed cleavage. **(F)** Sequence logos representing motifs surrounding non-targeted cleavage sites generated with the targeting and non-targeting guide RNAs. **(G)** CNNNG triplet heatmap detailing frequency of cleavage at each central triplet sequence across *λ* genome, normalized to genomic triplet abundance.Uncropped gel images for panels B-D are shown in Supplementary Fig. S10.

### Oxford Nanopore sequence mapping confirms CNNNG-dependence of non-targeted cleavage

To identify the NTCPs, we used Oxford Nanopore sequencing to map linear cleavage products from cleavage assays with *λ* DNA and TevCas12a/gRNA_*λ*_ or with TevCas12a/gRNA_AAVS1_ (Figs. 4). In each case, NTCPs were visible at 0.5 min and complete cleavage of the *λ* genome in to NTCPs by 8 min (Fig. 4B and 5D). We also observed NTCPs in the absence of a gRNA, consistent with data from cleavage of AAVS1 linear substrates. In experiments with TevCas12a and (gRNA_*λ*_), the on-target cleavage product mapped to the expected position (Fig. 4C). This product was not present in reactions with TevCas12a and gRNA_AAVS1_ (Fig. 4E). We sequenced reactions with Oxford Nanopore that were stopped after 2 minutes, and identified the Tev CNNNG cleavage motif from reads from experiments with the on-target gRNA_*λ*_ and non-targeted gRNA_AAVS1_ (Fig. 4F). Changing the NaCl concentration in the reaction buffer did not alter the Tev cleavage preference on *λ* DNA, as a highly similar CNNNG motif preference was observed under both low (12 mM) and high (102 mM) NaCl conditions (Fig. S5A-B). Moreover, we also identified a T/A preference at position 7 (2 bases downstream of the G of the CNNNG motif) (Fig. 4F) that is consistent with base preference for the wild-type I-TevI enzyme and other chimeric Tev-based nucleases^18^, ^31^. Using CNNNG sequences present in the mapped cleavage products, we determined that the preference for the NNN central triplet was very similar to that determined from previous studies (Fig. 4G), where GC-rich NNN triplets are cleaved less efficiently than AT-rich triplets^18^.

Further analysis of the Oxford Nanopore sequencing and mapping data revealed a relationship between the frequency of cleavage at CNNNG motifs and the distance to the nearest TTTV PAM sequence (Fig. 5). These findings indicate a decreasing activity on Tev CNNNG motifs as the spacing increases upstream from the TTTV PAM. This data is consistent with studies showing that the I-TevI nuclease displays a distance preference for CNNNG motifs spaced appropriately from the DNA-binding domain^32^. A similar distance relationship was found for fusion of the I-TevI nuclease to zinc fingers, TALE domains, or meganucleases^16–18^. We propose that scanning of DNA substrates for PAM sites by Cas12a and the subsequent steps for R-loop formation provide sufficient residence time for the Tev nuclease domain to cleave at CNNNG sites.

**Figure 5.**
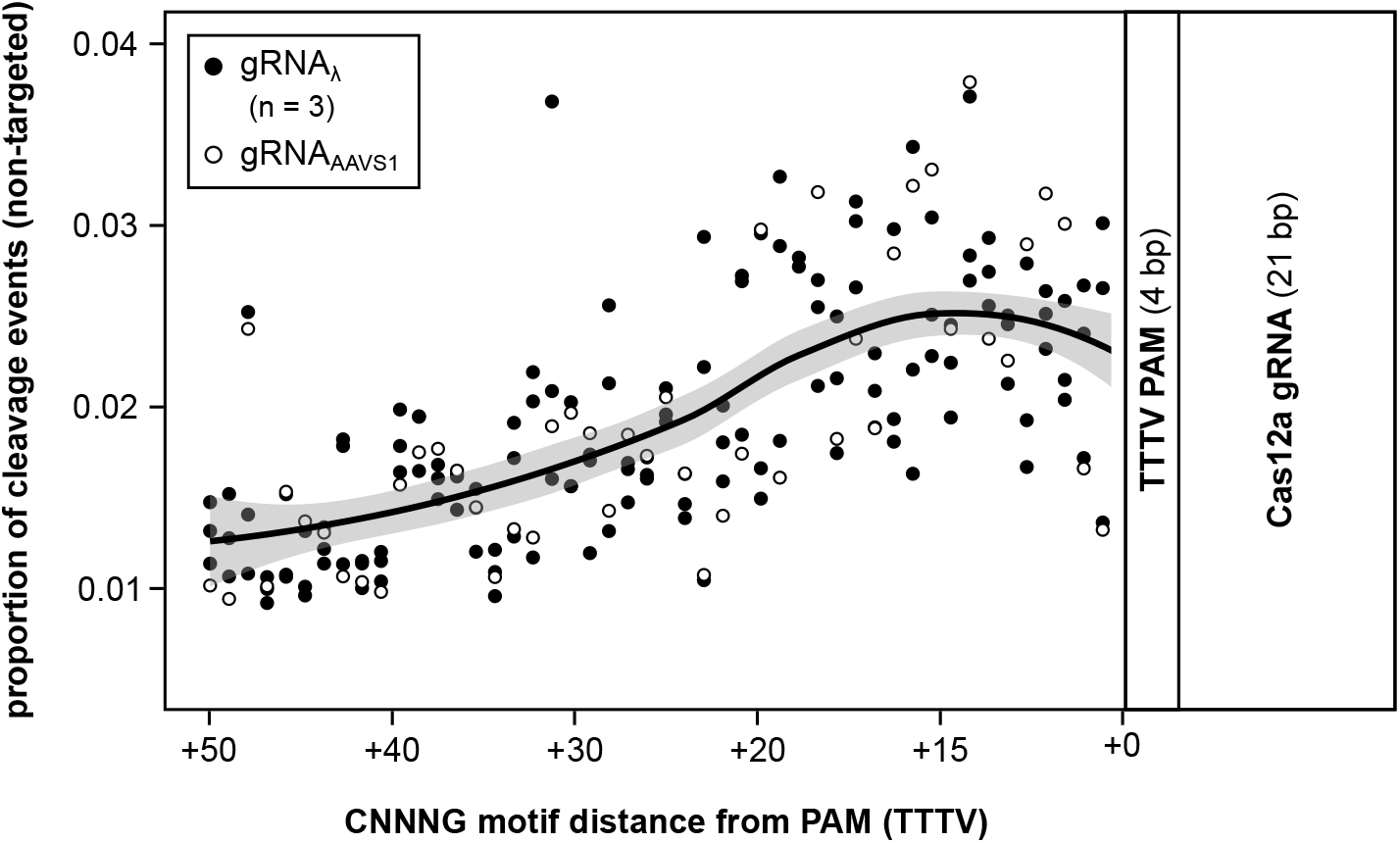
Cleavage of Tev CNNNG motifs decreases with distance from the TTTV PAM. Plot of the frequency of cleavage for mapped CNNNG motifs upstream (+ bp) of the TTTV PAM. Points are from four independent mapping experiments with targeted (gRNA_*λ*_) or non-targeted (gRNA_AAVS1_) guide RNAs. Shaded area indicates the 95% confidence interval for the fit of the data (non-parametric smoothing function).

### Cas12a and SaCas9 cleavage is sensitive to salt concentrations

It was recently shown that Cas12a searches for TTTV PAM sites by facilitated diffusion along a DNA substrate and that the rate of diffusion slows at lower salt concentrations^15^, ^33^. These observations suggest that Tev-directed NTCPs may result from Cas12a searching for TTTV PAM sites under the different salt concentrations used in the *in vitro* assays. To test this idea, we designed a 2000-bp linear “Cas-OFF” substrate that contained an overlapping, on-target AAVS1 Cas12a and SaCas9 site, as well as five sites that varied by one to three nucleotide transversion substitutions (Fig. 6A, S6A). The positions of the substitutions were PAM distal (*>*15 from the PAM), PAM medial (between 8-15 from the PAM), and PAM proximal (between 1-8 from the PAM) (Fig. 6A, S6A). Cleavage of each site by Cas12a would generate a discrete length product. When purified Cas12a/gRNA_AAVS1_ was incubated with the substrate across a 250-fold range of sodium chloride concentrations, we observed cleavage at each of the sites in the order on-target *>* PAM distal *>* PAM medial *>* PAM proximal (Fig. 6B-C). This cleavage preference is consistent with the tolerance of Cas12a/gRNA to target site mismatches^34^, ^35^.

**Figure 6.**
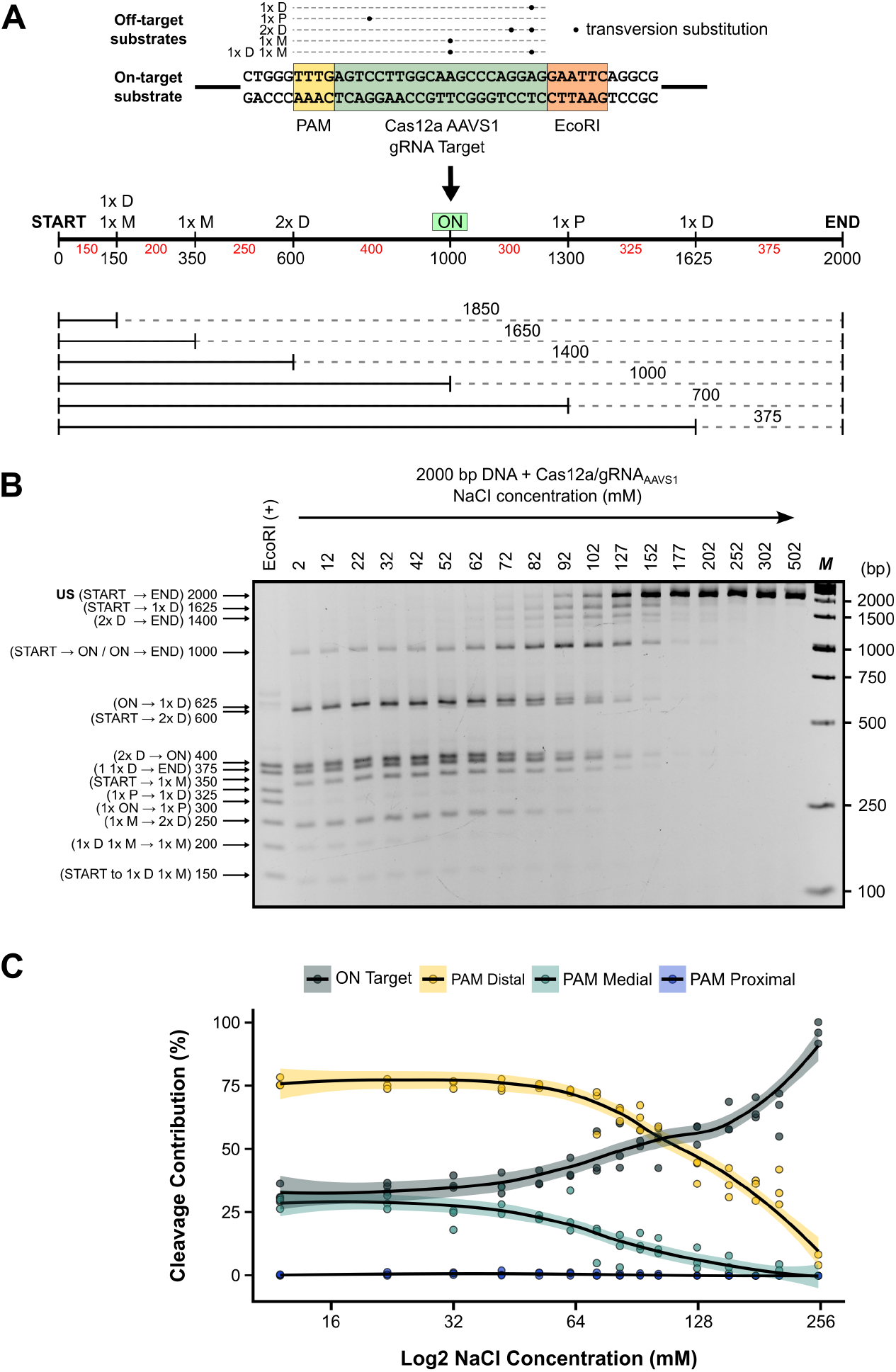
Cas12a cleavage at on-target and mismatched target sites is sensitive to salt concentration. **(A)** Schematic of the linear substrate with the on-target site flanked by an EcoRI site indicated. The position of mismatches to the on-target site are indicated above the sequence, and labeled by their position in the linear substrate shown below. The anticipated cleavage products and sizes are also indicated on the linear substrate. **(B)** Representative agarose gel of Cas12a cleavage under the indicated NaCl concentrations. Sizes and identity of products are indicated on the left of the gel. US, uncut substrate. **(C)** Plot of cleavage of on-target or mismatched target sites grouped by the position of the mismatches. Points are three independent replicates with the shaded area indicated the 95% confidence interval for fit of the data to a non-parametric smoothing function. Uncropped gel image for panels B is shown in Supplementary Fig. S11.

We performed a similar experiment with SaCas9, titrating salt concentration in a reaction containing 1 *×* CutSmart and a substrate with on-target and mis-matched target sites (Fig. S6B-C). Similar to that observed with Cas12a, SaCas9 displayed differential cleavage activity at the mismatched target sites relative to the on-target site at salt concentrations below 128 mM. At concentrations above 128 mM, increased fidelity towards on-target cleavage was observed.

### Non-targeted cleavage by TevSaCas9, Tev-ZFE and MegaTevs

We were also interested in determining if the NTCPs were a unique feature of the TevCas12a architecture, or whether cleavage activities of other Tev-based editors could be modulated by changing *in vitro* salt concentrations. To this end, we used previously constructed Tev-gene editors; MegaTevs, a fusion of Tev to the I-OnuI meganuclease^18^; Tev-ZFE, a fusion of Tev to the RyA zinc finger^16^; and TevSaCas9, a Tev fusion to the Cas9 nuclease from *Staphylococcus aureus*^20^ (Fig. 7A-C). To target TevSaCas9, we synthesized an SaCas9 gRNA that targeted the *λ N* gene. When *λ* DNA was incubated with the Tev-ZFE, MegaTev or TevSaCas9/gRNA_*λ*_ in buffer containing 50 mM sodium chloride and 1 *×* rCutSmart buffer, NTCPs were observed for all three fusions (Fig. 7D-F). A product consistent with gRNA-directed cleavage by the TevSaCas9 fusion was visible (cleavage product, CP) (Fig. 7F). No discrete, targeted cleavage products were visible in reactions with the MegaTev or Tev-ZFE fusions (Fig. 7D and 7E), consistent with the fact that the meganuclease and RyA zinc finger component of the fusions were not engineered to cleave the *λ* genome. Mapping the cleavage products for the three editors using Oxford Nanopore revealed a consensus Tev cleavage motif for all three fusions (Fig. 7D-F).

**Figure 7.**
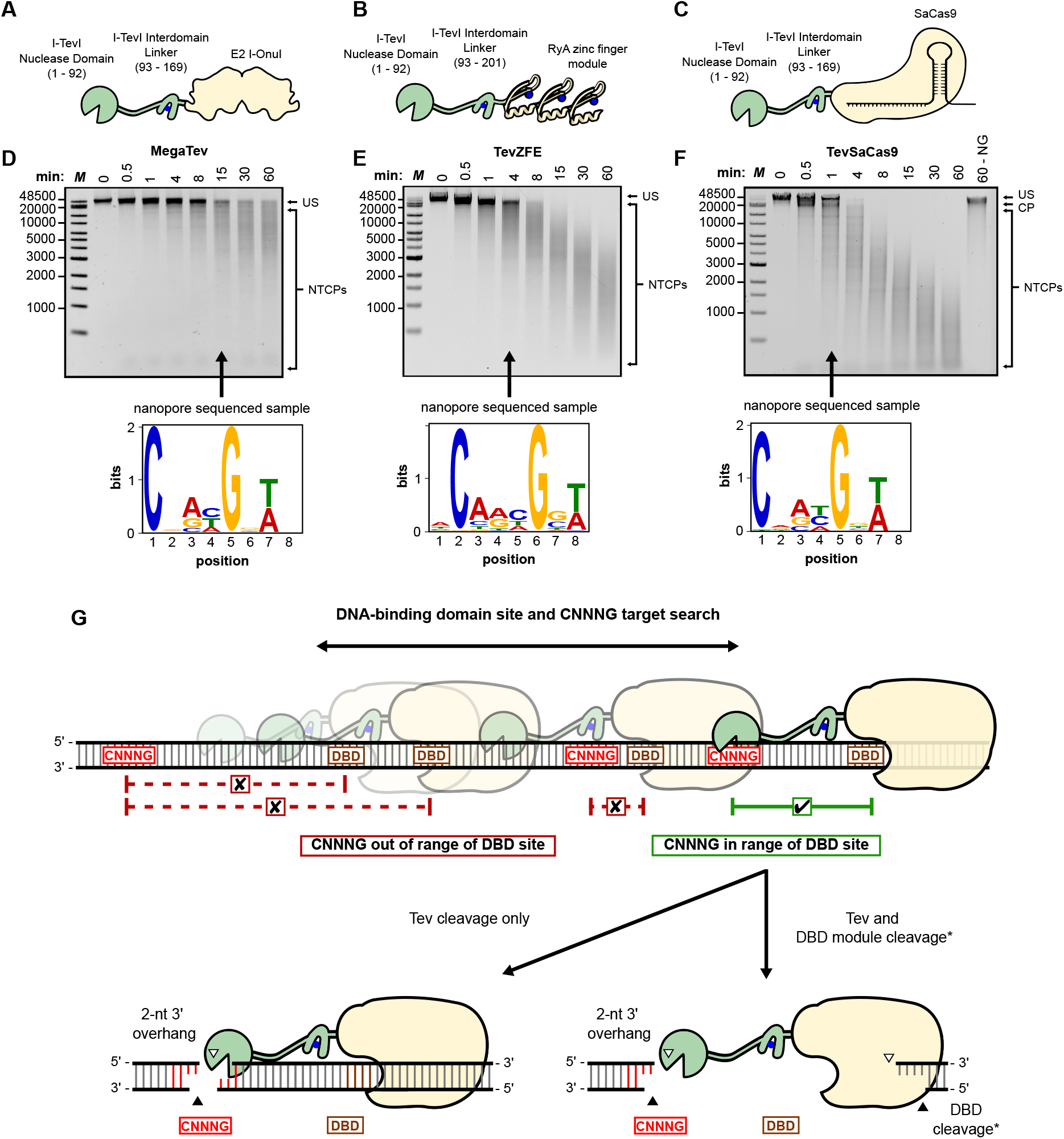
Non-targeted cleavage products are observed with other chimeric gene editors. **(A)-(C)** Shown are schematics of Tev nuclease domain fusions to different DNA-binding domains and **(D)-(F)** representative time-course cleavage assays of each fusion with *λ* DNA. The time point used for Oxford Nanopore sequencing is indicated. The sequence logos identified for each I-TevI fusion are shown below each gel image. US; unreacted substrate; NTCPs; non-targeted cleavage products. **(G)** Model of non-targeted cleavage by chimeric Tev-based nucleases. Tev editors engage with DNA substrate and scan for target sites (DBD). If engaged at at a DBD and a Tev CNNNG motif is in range (green), cleavage occurs at the Tev and DBD module sites (in the case of active TevCas fusion proteins). Cleavage by Tev does not proceed if Tev editors are engaged at DBD sites where the CNNNG motif is out of range (red). Uncropped gel images for panels D-F are shown in Supplementary Fig. S12.

Collectively, this data shows that the activity and specificity of gene editors that use RNA- and protein-based targeting mechanisms can be modulated *in vitro* by simply changing the buffer salt concentrations. Lower salt conditions promote stability of the chimeric editors at sub-optimal target sites, enabling Tev cleavage at appropriately positioned CNNNG motifs (Fig. 7G). The cleavage preference of the Tev nuclease domain does not change regardless of the fusion context. This indicates that the effect of lowering the salt concentration is to change the stability of the Tev domain on DNA, rather than to change or relax sequence preference.

## Discussion

Our *in vitro* characterization of TevCas12a activity revealed an unanticipated result; changing buffer salt concentrations effectively decoupled Tev cleavage from Cas12a-gRNA targeting and resulted in the appearance of non-gRNA directed cleavage that mapped to Tev CNNNG motifs. This decoupling was also observed for other Tev-based editors that use RNA- and protein-based targeting mechanisms. Beyond the immediate biochemical findings, de-coupling of cleavage activity from DNA targeting provides a straightforward workflow to profile the cleavage preference and specificity of nuclease domains with low DNA-binding affinity. This is particularly relevant for the GIY-YIG superfamily that is vastly underexplored in terms of cleavage preferences: of the many predicted GIY-YIG nuclease domains, only I-TevI and I-BmoI have been rigorously characterized^31, 36–38^. The ability to tether stand-alone nuclease domains to a DNA-binding module like Cas12a enables interrogation of cleavage preferences in a systematic and high-throughput fashion. Because the Cas12a scaffold can localize the fusion protein to DNA substrates without requiring sequence-specific targeting, it serves as a biochemical “anchor” to position weak-binding nuclease domains at a variety of diverse targets, providing a unique opportunity to expand the functional sequence space of GIY-YIG domains, or other nucleases.

GIY-YIG nucleases are compact modular domains that are found in a variety of protein scaffolds^30, 39–41^. The Tev nuclease domain (minimally residues 1-92) has very low affinity for DNA and an exact determination of binding affinity has been hindered by the fact that the wild-type nuclease domain (or full-length I-TevI) is extremely toxic in *E. coli* and recalcitrant to cloning and overexpression^29^, ^42^. Studies with the nuclease domain from the isoschizomer I-BmoI revealed a binding affinity of 10 µM; a longer construct, encompassing residues 1–154 and analogous to the Tev 1–169 construct used here, had an affinity of 500 nM^37^. While these measurements—and the 1.1 nM dsDNA binding constant reported for Cas12a^27^— were obtained at different salt concentrations, they nonetheless highlight the much weaker affinity of the Tev nuclease domain. Thus, our data indicate that non-targeted cleavage products observed with TevCas12a (and other editors) arise through DNA binding domain-mediated DNA interactions. While PAM recognition enhances cleavage at proximal CNNNG motifs, transient or non-canonical DNA contacts by Cas12a are still sufficient to stabilize Tev binding and enable cleavage at more distal CNNNG sites. Indeed, previous studies with Tev-zinc finger, Tev-meaganuclease, and Tev-SpCas9 fusions showed that Tev cleavage was dependent on targeted DNA-binding^16, 18^, ^19^. Moreover, when expressed as a component of dual-nuclease editors such as MegaTev or TevCas9, the Tev nuclease domain has not displayed toxicity in either *E. coli* or human cells, as would be expected if Tev were non-specifically interacting with and cleaving DNA^18^.

The sensitivity of protein-DNA interactions to *in vitro* buffer conditions is well known.^7–9^. For instance, the rate of non-specific cleavage by FokI-based zinc finger nucleases can be enhanced in low salt buffers^10^, and the interactions of SpCas9 with non-specific sites is not impacted by changes in salt concentration^13^, ^14^. For TevCas12a, low sodium chloride concentrations resulted in the appearance of cleavage products in 1 *×* CutSmart-buffered reactions not directed by Cas12a/gRNA binding. This de-coupling of DNA cleavage from gRNA targeting is consistent with studies showing that Cas12a scans for TTTV PAM sites by facilitated diffusion, and that the rate of diffusion slows at lower salt concentrations^15^, ^33^. We found that non-targeted cleavage products are enhanced in low salt buffer, consistent with reduced *K*_*off*_ values that stabilize Tev–DNA complexes so that *K*_*cat*_ and *K*_*off*_ are of similar magnitude, enabling productive cleavage. In this context, Cas12a samples DNA for PAM sites through facilitated diffusion, in which sliding and hopping events along DNA increase the efficiency of target search relative to three-dimensional collisions driven by Brownian motion alone, extending local residence time for Tev cleavage at CNNNG motifs. Under low-salt conditions, reduced ionic competition at the DNA interface (‘ionic thermodynamic buffering’) stabilizes weak protein–DNA interactions, extending the dwell time of Cas12a–Tev complexes and promoting Tev cleavage at CNNNG motifs not strictly coupled to Cas12a gRNA targeting. The positional dependence of CNNNG cleavage sites within a defined window upstream of TTTV PAMs also supports the idea that Tev cleavage is linked to diffusion of Cas12a along substrate. Tev non-targeted cleavage products are reduced at *>* 100 mM sodium chloride, presumably due to a faster facilitated Cas12a diffusion rate, decreased ionic thermodynamic buffering at sub-optimal Cas12a sites, and the inherently weak DNA affinity of the Tev nuclease domain. Given that all reactions contained an rCutSmart contribution of 50 mM K^+^, these conditions correspond to total monovalent cation concentrations above 150 mM. Low (12 mM) or high (102 mM) NaCl buffer did not change Tev cleavage preference; rather, it only modulates the cleavage kinetics. The fact that CNNNG motif preference remains conserved across different fusion partners—Cas12a, Cas9, meganucleases, and zinc fingers—underscores the biochemical autonomy of the Tev domain and mirrors the modularity of GIY-YIG homing endonucleases observed in nature. This modular behavior reinforces the concept that GIY-YIG domains can be re-deployed in new architectures while retaining intrinsic biochemical features, supporting their further exploration as versatile genome editing components.

Although the primary motivation of this study was to develop *in vitro* methodologies to map the preference for Tev cleavage, a question arising from our study is the *in vivo* relevance of non-targeted cleavage by Tev-based chimeric nucleases. In this regard, TevSpCas9 cleaves at targeted sites in mam-malian cell lines^19^, ^43^ and in diatoms^44^, ^45^. Cas12a and SaCas9 also cleave at targeted sites in cells and are not cytotoxic. The tolerance of Cas12a and SaCas9 to target DNA mismatches in low salt conditions is consistent with stabilization of protein-DNA (or RNA-DNA) interactions that, in the context of a chimeric editor, facilitate Tev interrogation for CNNNG motifs. The opposing idea, that the Tev nuclease domain cleaves non-specifically and independently of DNA-binding domain tethering at CNNNG motifs, is not supported by the biochemical properties of the Tev nuclease domain or the observed activities of Tev-based chimeric editors where targeting is a function of the DNA-binding domain. Our data point to a remarkably consistent and specific preference of the Tev nuclease for CNNNG motifs across different fusion contexts and buffer conditions.

In summary, we have shown that changes to salt concentrations *in vitro* can modulate the cleavage activity of chimeric gene editors to de-couple nuclease domain cleavage from DNA targeting. This effect was seen for editors that use RNA- or DNA-based targeting mechanisms. From a practical standpoint, our data suggest that *in vitro* buffer conditions should be carefully considered when profiling the specificity of editors based on CRISPR nucleases, as slight changes in salt concentration (or other buffer components) could impact the activity and the readout of the tolerance to mismatches in the target site. In theory, this approach could profile the preference of any nuclease fused to a targeting domain^46^, ^47^, or to determine the sequence preferences of other DNA modifying domains, including base editors^48^ or methylases^49^. This approach would also enable mutational dissection of the Tev nuclease domain and linker to explore how specific residues contribute to cleavage-site selection, motif recognition, or catalytic activity, potentially informing the design of synthetic nucleases with tailored specificity.

## Materials and Methods

### Bacterial strains

*Escherichia coli* NEB5*α* F’I’ (F’ *proA+B+ lacI*^*q*^ *δ(lacZ)M15 zzf::Tn10 (TetR)* /*fhuA2δ(argF-lacZ)U169 phoA glnV44 φ* 80*δ(lacZ)M15 gyrA96 recA1 relA1 endA1 thi-1 hsdR17*) and *E. coli* ER2566 *F*^*-*^ *mcrA* (*mrr-hsdRMS-mcrBC*) 80*lacZ M15 lacX74 recA1 ara139 (ara-leu)7697 galU galK rpsL* (*StrR*) *endA1 nupG* (NEB) were used for cloning and protein expression and were grown in LB media.

### Plasmid construction

All TevCas12a variants were cloned into the pET11a expression vector using Gibson Assembly with NEBuilder® HiFi, following the manufacturer’s protocol. To create TevCas12a fusions, *E. coli* codon-optimized I-TevI (amino acids 1–169, GenBank: P13299.2) with a GGSGGTGGSG peptide at the C-terminal end was fused to the N-terminus of AsCpf1 (Cas12a, GenBank: U2UMQ6.1). Mutations that inactivated I-TevI (R27A) or Cas12a (D908P) were introduced into the wild-type TevCas12a plasmid using golden gate mutagenesis. Primers for mutagenesis and cloning are listed in Table S1. All constructs were verified by Sanger sequencing of the cloned insert (London Regional Genomics Facility, London, ON) or whole plasmid sequencing (Plasmidsaurus). A list of oligonucleotides used in cloning and plasmids used in this study are provided in Table S1 and S2, respectively.

### Guide RNA synthesis

All guide RNA sequences and oligonucleotides are listed in Table S1. Guide RNAs were synthesized by one-pot *in vitro* transcription using the T7 RiboMAX^®^ Express Large Scale RNA Production System (Promega). DNA templates were prepared using two distinct strategies depending on the nuclease. For Cas12a gRNAs, templates were formed by annealing fully complementary sense and antisense oligonucleotides (2 *µ*M each; Integrated DNA Technologies), yielding a 69 bp duplex encoding a T7 promoter, spacer, gRNA scaffold, and a 3’ poly-T terminator. No template extension was required. For SaCas9 gRNAs, templates were generated by annealing a guide-specific forward oligo (containing the T7 promoter, spacer, and partial scaffold) with a universal reverse oligo (DE6018) encoding the remainder of the scaffold and a T-rich terminator. These partially overlapping oligos were extended in situ using Klenow Fragment (3’ → 5’ exo-, New England Biolabs). Final 20 *µ*L reactions contained 2.5 U Klenow, 125 *µ*M dNTPs, 1× RiboMAX Express T7 Buffer, 2 *µ*L T7 Express Enzyme Mix (Promega), and the annealed DNA duplex. Reactions were incubated at 37^*°*^C for 4 h. Transcription products were purified using the Monarch^®^ RNA Cleanup Kit (New England Biolabs) according to the manufacturer’s protocol. No DNase treatment was performed. RNA integrity was assessed by agarose gel electrophoresis.

### DNA substrates for *in vitro* assays

AAVS1 substrates were amplified from HEK293 genomic DNA using PrimeSTAR GXL DNA polymerase (Takara Bio) and primers OL0345 and OL0346. The 872 bp amplicon corresponds to the AAVS1 locus (GenBank: AC010327.8) and was cloned into the pJET vector (pJET-AAVS1) using the ThermoFisher CloneJET PCR Cloning Kit. Sequence integrity was confirmed by Sanger sequencing of the genomic PCR product, and again following cloning.

A linear Cas9/Cas12a off-target DNA substrate (“Cas-OFF”) was assembled by BsaI-HFv2 Golden Gate cloning of five gBlocks (Integrated DNA Technologies), each flanked by BsaI recognition sites. Equimolar amounts (0.05 pmol) were mixed and ligated using BsaI-HFv2 (New England Biolabs), T4 DNA Ligase (New England Biolabs), and 1 *×* T4 DNA ligase buffer (New England Biolabs). The assembled product was used directly as template for PCR amplification with primers DE7182 and DE7184.

Substrates lacking CNNNG and TTTV motifs were PCR-amplified from gBlocks (IDT) using gBlocks as templates with primers DE6502 through DE6785. Lambda DNA was purchased from New England Biolabs.

### Protein purification

C-terminal 6×His-tagged TevCas12a variants and TevSaCas9 (in pET11a), as well as MegaTev and TevRyA(ZFE) (in pACYC-Duet1), were expressed under the control of a T7 promoter in *E. coli* ER2566 cells. Cultures (250 mL) were grown in a 2:1 flask-to-volume ratio at 37°C to an OD_600_ of 0.6, then induced with 1 mM IPTG and incubated overnight at 16°C.

Cells were harvested by centrifugation and lysed by sonication in binding buffer [20 mM Tris*·*HCl, pH 8.0, 500 mM NaCl, 1 mM DTT, 5% (v/v) glycerol, 10 mM imidazole] supplemented with 1 mM phenylmethylsulfonyl fluoride (PMSF). Clarified lysates were applied to 1 mL HisTrap™ Ni-NTA affinity columns (Cytiva), and bound proteins were eluted in a stepwise gradient of imidazole (100, 200, 300, and 500 mM). Eluted fractions were dialyzed using 8 kDa MWCO tubing into storage buffer (identical to binding buffer but lacking imidazole), aliquoted, flash-frozen in a –80°C ethanol bath, and stored at –80°C.

### *In vitro* cleavage reactions

Cleavage reactions were performed in biological triplicate at varying molar ratios of RNP to substrate (20:1, 10:1, 5:1, 1:1, or 1:5) using TevCas12a RNPs, TevSaCas9 RNPs, MegaTev, or TevRyA(ZFE). RNPs were assembled at a 1:1.5 molar ratio of protein to guide RNA by incubation at room temperature for 10 minutes. Reactions contained 2.5 nM DNA substrate in 50 mM NaCl, 20 mM Tris *·* HCl (pH 8.0), 1 mM DTT, and 1 *×* rCutSmart buffer (New England Biolabs; 50 mM potassium acetate, 20 mM Tris-acetate, 10 mM magnesium acetate, 100 *µ*g/mL BSA), and were incubated at 18°C, 24°C, 30°C, or 37°C. Aliquots were taken at designated time points and quenched in stop solution (16.7 mM EDTA, 50 *µ*g/mL RNase A, 50 *µ*g/mL proteinase K), followed by incubation at 37°C for 30 minutes.

For salt titration experiments, TevCas12a, Cas12a and SaCas9 RNPs were assembled and diluted to 500 nM in nuclease-free water to minimize salt carryover from the storage buffer. NaCl was titrated (0–500 mM final) into reaction mixtures containing 2.5 nM *λ* DNA or a 2000 bp linear “Cas-OFF” amplicon in 20 mM Tris *·* HCl (pH 8.0), 1 mM DTT, and 1 *×* rCutSmart buffer. RNPs were added last at a 20:1 RNP:DNA molar ratio, and reactions were incubated at 37°C for 1 hour (TevCas12a) or 15 minutes (Cas12a, SaCas9) before quenching.

Cleavage products were analyzed by electrophoresis on 12% (v/v) polyacrylamide gels in Tris-borate-EDTA (TBE) buffer for linear substrates, or 0.8% (w/v) agarose gels in Tris-acetate-EDTA (TAE) buffer for pJET-AAVS1 and *λ* DNA. Gel images were analyzed in ImageLab (BioRad) by manual band calling based on fragment size, and cleavage fractions were determined from relative band intensities.

### Oxford Nanopore sequencing and data analysis

Pooled cleavage reactions containing 1.5 *µ*g of pJET-AAVS1 or *λ* DNA were purified using the Monarch PCR & DNA Cleanup Kit (New England Biolabs) according to the manufacturer’s protocol. End-repair and dA-tailing were performed using the NEBNext^®^ Companion Module, followed by ligation of bar-coded adapters from the Native Barcoding Expansion kit (EXP-NB104, Oxford Nanopore Technologies). Ligations were incubated at room temperature for 30 minutes and purified using AMPure XP beads. A total of 800 ng of barcoded product was recovered and split equally across samples. Final pooled input for sequencing was 500 ng. Library preparation followed the SQK-LSK114 protocol. DNA concentration and integrity were assessed using a Qubit fluorometer and agarose gel electrophoresis.

Sequencing was performed on MinION flow cells (R9.4.1), yielding 2–3 million reads per run. Basecalling and demultiplexing were performed with Guppy (v6.5.7) using default parameters. Reads were aligned to the corresponding reference sequences (pJET-AAVS1 and *λ* DNA) using minimap2 with the -ax map-ont preset. Mapped reads were parsed using a custom Perl script to extract read length, strand orientation, and mapped position. Cleavage sites were defined as the mapped termini of reads relative to the reference sequence.

Custom Python scripts were used to identify cleavage positions and characterize sequence features at, upstream, and downstream of each site. Cleavage frequencies by position were calculated, and recurrent motifs and PAM proximities were identified. Motif logos were generated using MEME (v5.5.5) with the “one occurrence per sequence” (oops) model.

To assess proximity to protospacer-adjacent motifs, each cleavage site was scanned for the nearest upstream TTTV [0] PAM (defined by end coordinate and strand mapping), and the nearest downstream or BAAA [16] PAM (defined by start coordinate and strand mapping). Linear distances between cleavage positions and PAM sites were calculated. All analyses were performed using custom Python scripts provided in Supplementary Information.

## Acknowledgements (not compulsory)

The authors thank members of the Edgell lab and Specific Biologics Inc. for comments on the manuscript.

## Author contributions statement

K.W.L., A.L.W, T.A.M., B.E.S. and D.R.E conceived the experiments. K.W.L., T.A.M., B.E.S. and A.L.W conducted the experiments. K.W.L., A.L.W and D.R.E analyzed the results. K.W.L. and D.R.E. wrote the manuscript. K.W.L., A.L.W, T.A.M., B.E.S. and D.R.E read the manuscript.

## Data availability

The Oxford Nanopore sequences generated in this study have been deposited in the Sequence Read Archive [PRJNA1273990]. Computer code is provided in the Supplementary Information.

