## Supplemental Data for "A buffer-tuning strategy to profile domain-specific activity of chimeric I-TevI/CRISPR gene editors *in vitro*"

Loedige et. al

### Supplementary Figures

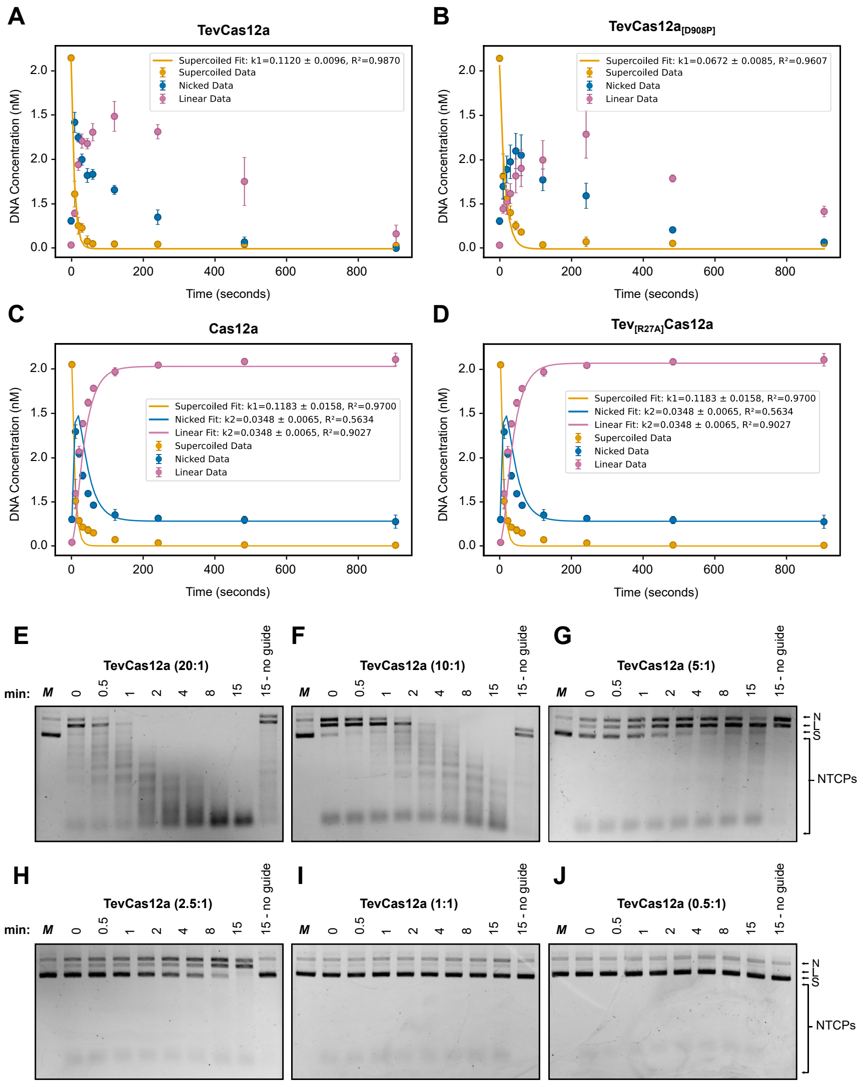


Figure S1: Kinetic analysis of TevCas12a and enzyme turnover under DNA excess. (A–D) Cleavage kinetics of supercoiled plasmid DNA by TevCas12a variants. Supercoiled (yellow), nicked (blue), and linear (pink) species were quantified and fit to exponential models. (E–J) Visualization of cleavage timecourse gels of TevCas12a cleavage with decreasing protein:DNA ratios (E: 20:1 to J: 0.5:1). Efficient non-targeted cleavage is lost at low enzyme excess, consistent with single-turnover activity. NTCPs = non-targeted cleavage products; N = nicked; L = linear.

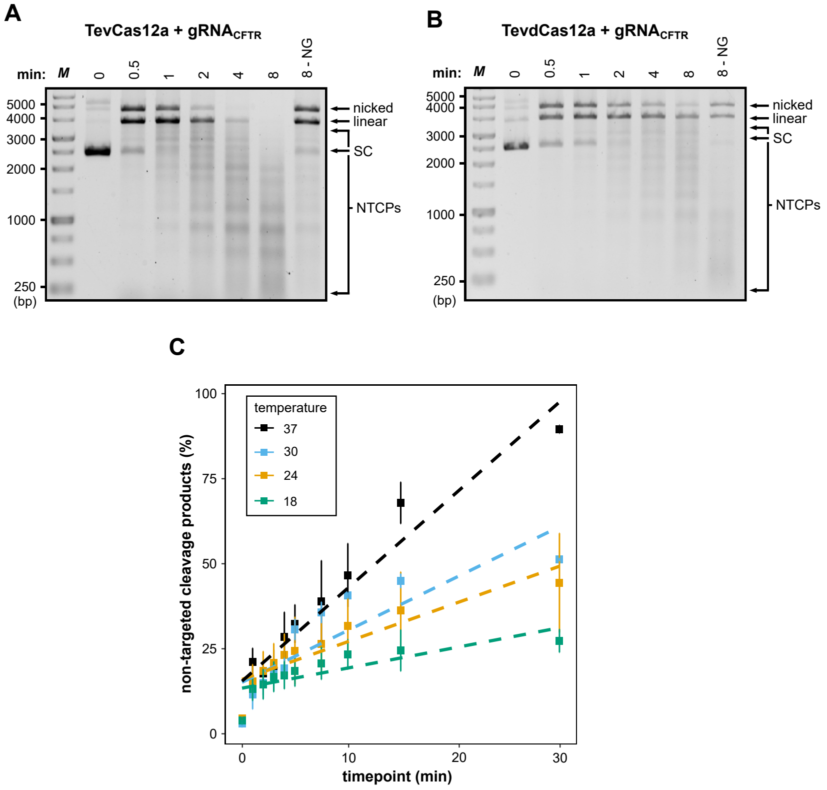


Figure S2: Non-targeted cleavage by TevCas12a is Tev-dependent, guide RNA–independent, and temperature-dependent. (A-B) Cleavage of plasmid by TevCas12a or TevdCas12a with non-targeted $gRNA_{CFTR}$ displays non-targeted cleavage. SC, supercoiled substrate. NTCPs, non-targeted cleavage products. (C) Quantification of non-targeted cleavage products over time at different temperatures. Non-targeted cleavage is minimal at 18°C and maximal at 37°C.

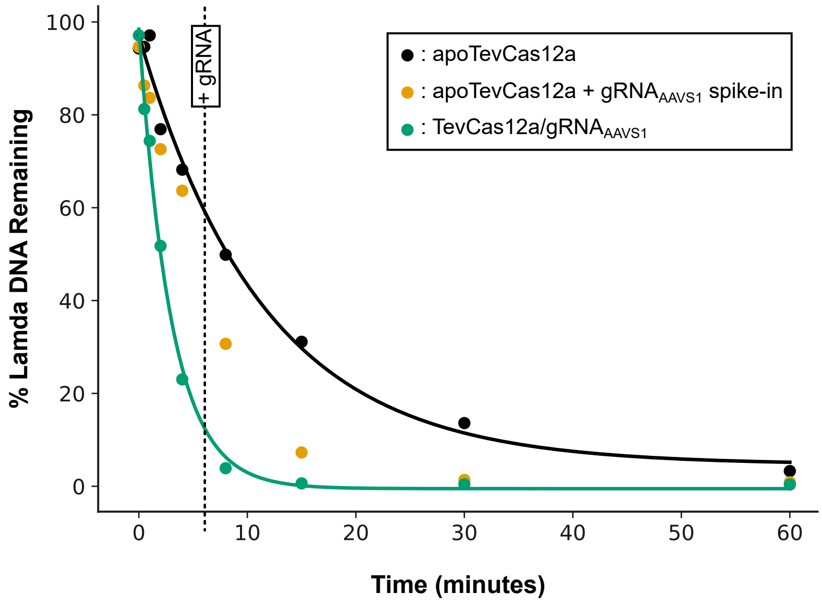


Figure S3: Quantification of NTCP formation under different stimulatory gRNA conditions. Percent $\lambda$ DNA remaining over time was quantified from agarose gels shown in Fig. 2C (apo-TevCas12a), Fig. 2D (apo-TevCas12a with gRNA$_{AAVS1}$ spiked in at 8 min), and Fig. 4D (TevCas12a/gRNA$_{AAVS1}$). Both spiked and non-targeted gRNAs stimulated NTCP formation relative to apo-TevCas12a. Both apo-TevCas12a and reactions initiated with gRNA$_{AAVS1}$ followed exponential decay kinetics of $\lambda$ substrate, whereas the mid-reaction gRNA spike-in deviated from this trajectory by accelerating NTCP formation.

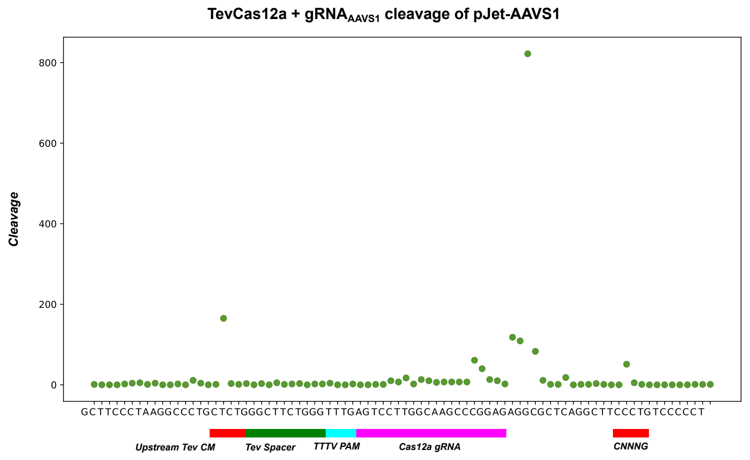


Figure S4: Oxford Nanopore mapping of TevCas12a cleavage product abundance at on-target loci. Cleavage profile of TevCas12a with gRNA$_{AAVS1}$ on the pJET-AAVS1 plasmid. Each point represents total cleavage events by position near the annotated AAVS1 target site. Cleavage is observed at Cas12a and Tev-specific target motifs, consistent with dual-nuclease activity.

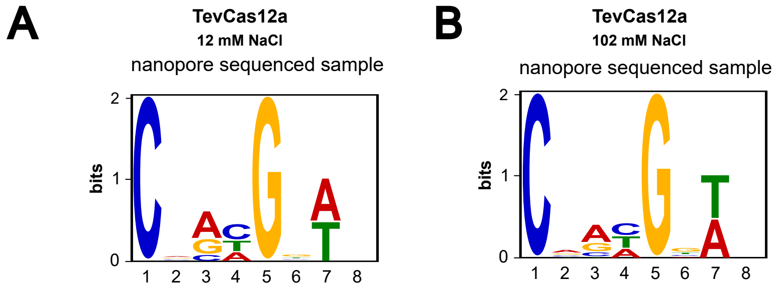


Figure S5: Effect of salt concentration on TevCas12a cleavage activity and preference. (A–B) Sequence logos of cleavage sites derived from nanopore sequencing of products generated by TevCas12a at 12 mM (A) and 102 mM (B) NaCl. Cleavage preference identity across salt concentration, indicates unaltered sequence specificity.

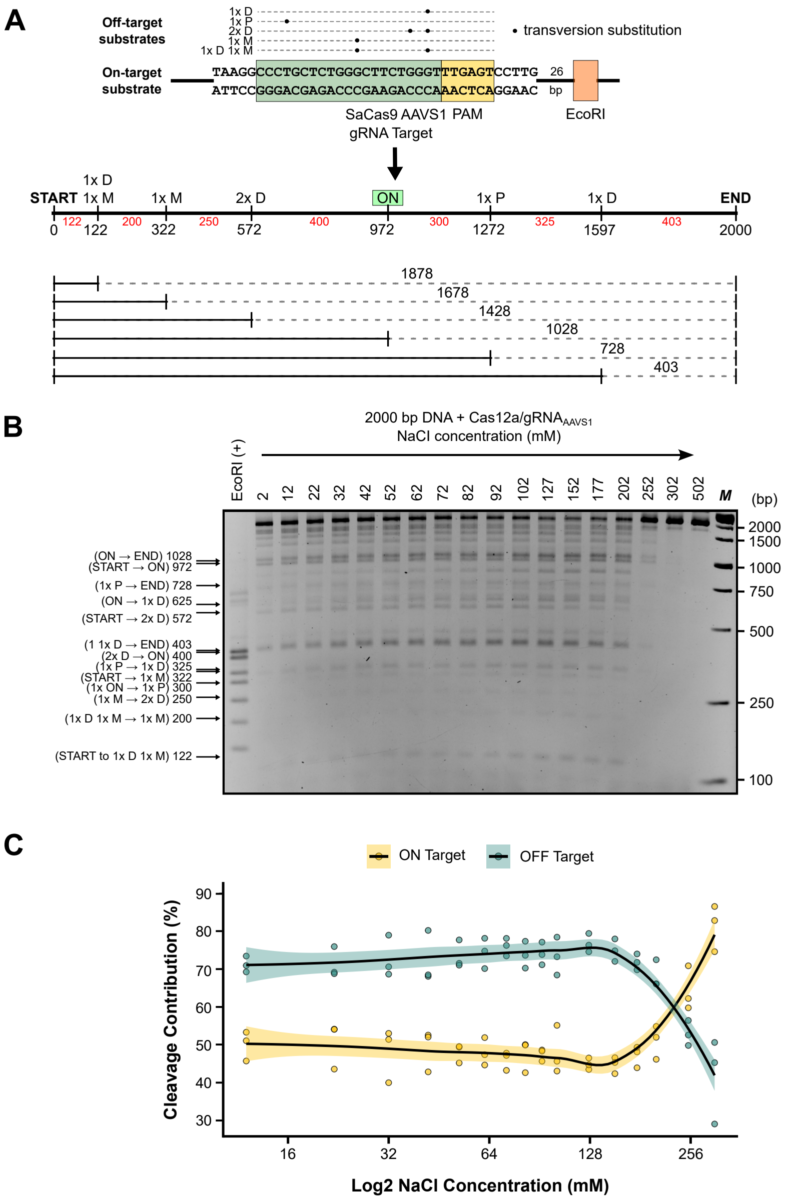


Figure S6: SaCas9 cleavage at on-target and mismatched target sites is modulated by salt concentration. (A) Schematic of the linear DNA substrate containing an on-target site for SaCas9/gRNA$_{AAVS1}$ (green) and a panel of mismatched off-target substrates with single or double transversion substitutions. Positions of mismatches and expected cleavage products are annotated relative to the 2000 bp linear substrate. (B) Representative agarose gel showing cleavage products generated by SaCas9 across a range of NaCl concentrations. Product sizes and identities are indicated on the left. (C) Quantification of cleavage at on-target (yellow) and mismatched off-target (blue) sites across salt concentrations. Data represent three independent replicates; shaded regions indicate 95% confidence intervals for nonlinear regression fits. At higher salt, cleavage shifts in favor of the on-target site.

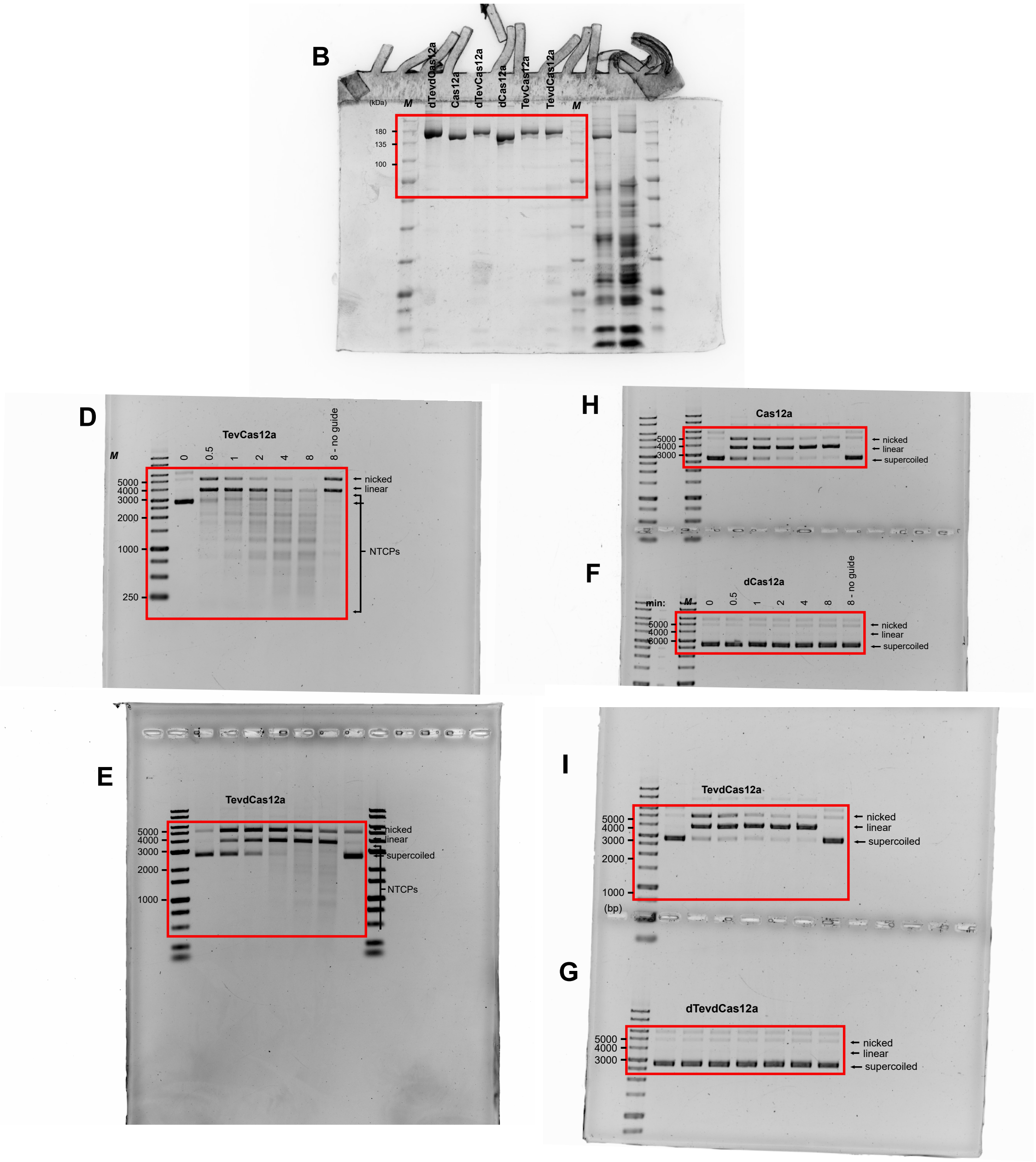


Figure S7: Uncropped agarose and SDS–PAGE gels corresponding to main text **Fig. 1 panels B–I**. Full gel images are shown for the purified TevCas12a and Cas12a variants (panel B) and for time-course cleavage assays of pJET-AAVS1 with TevCas12a/gRNA$_{\mathrm{AAVS1}}$ and catalytic mutants (panels D–I). Lanes correspond exactly to those in the main figure; molecular-weight (kDa) or size (bp) markers (M) are indicated for reference.

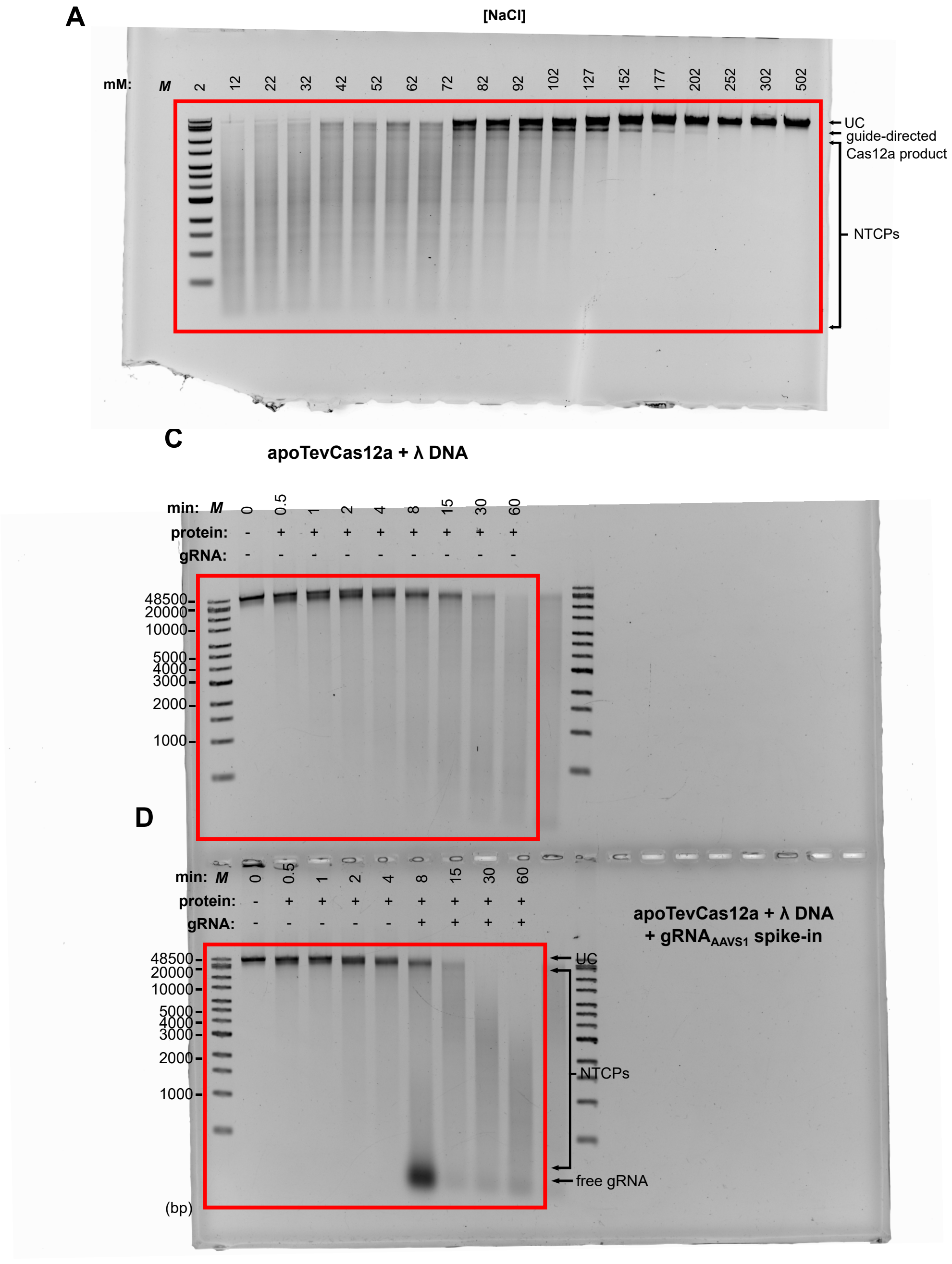


Figure S8: Uncropped agarose gels corresponding to main text **Fig. 2 A–D**. Full gel views for NaCl titration assays of TevCas12a/gRNA$_\lambda$ cleavage of $\lambda$ DNA (panel A), apo-TevCas12a time-course reactions (panel C), and gRNA$_{\mathrm{AAVS1}}$ spike-in assays (panel D). Sodium-chloride concentrations or reaction times are labeled as in the main text for reference. UC = uncleaved substrate; NTCPs = non-targeted cleavage products.

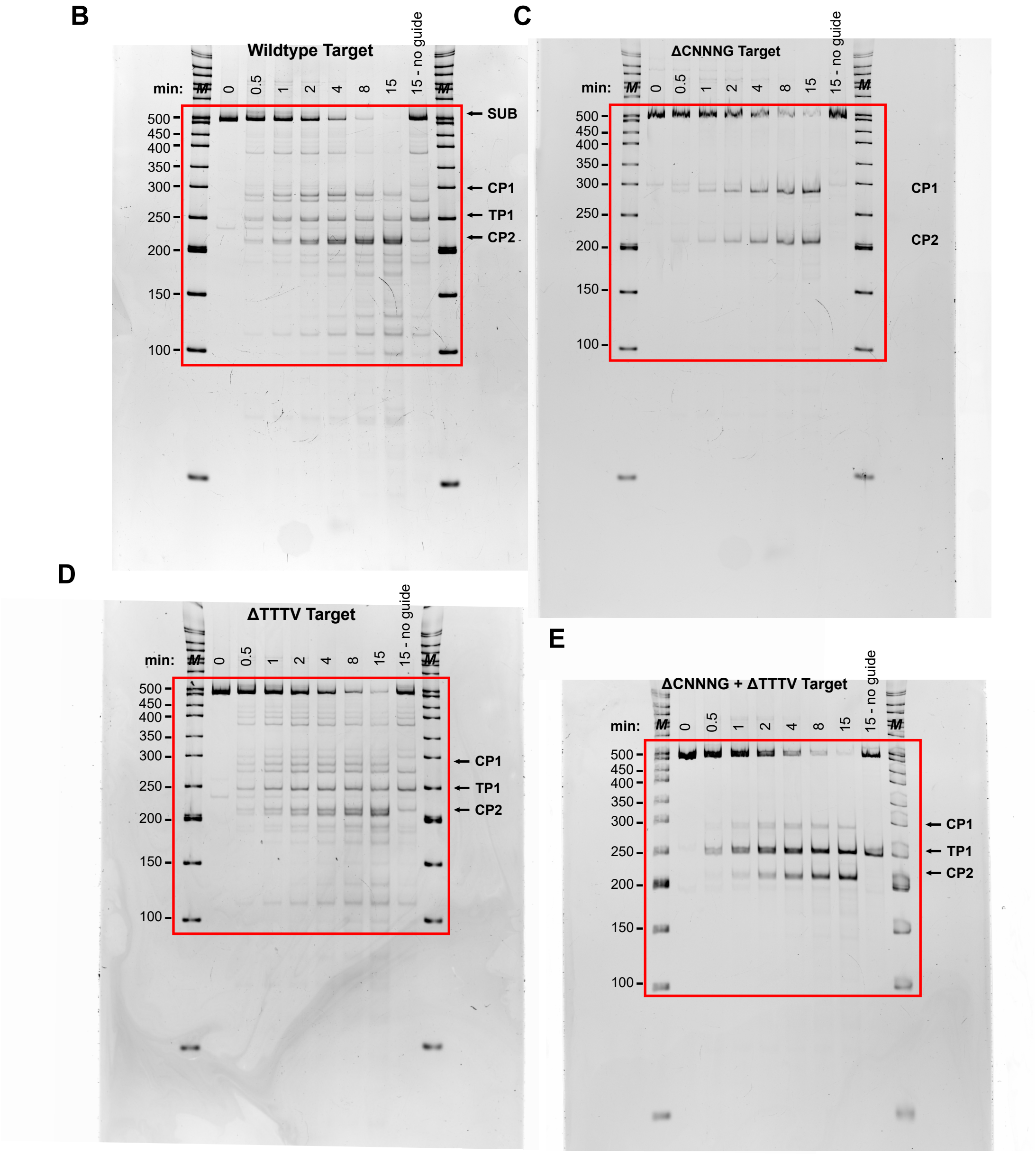


Figure S9: Uncropped polyacrylamide gels corresponding to main text **Fig. 3 B–E**. Full images of cleavage time-courses of wild-type and motif-mutated AAVS1 substrates (ΔCNNNG, ΔTTTV, ΔCNNNG + ΔTTTV). Lanes and time points match the cropped panels shown in the main figure for reference. SUB = uncleaved substrate; TP1 = Tev product; CP1–2 = Cas12a cleavage products.

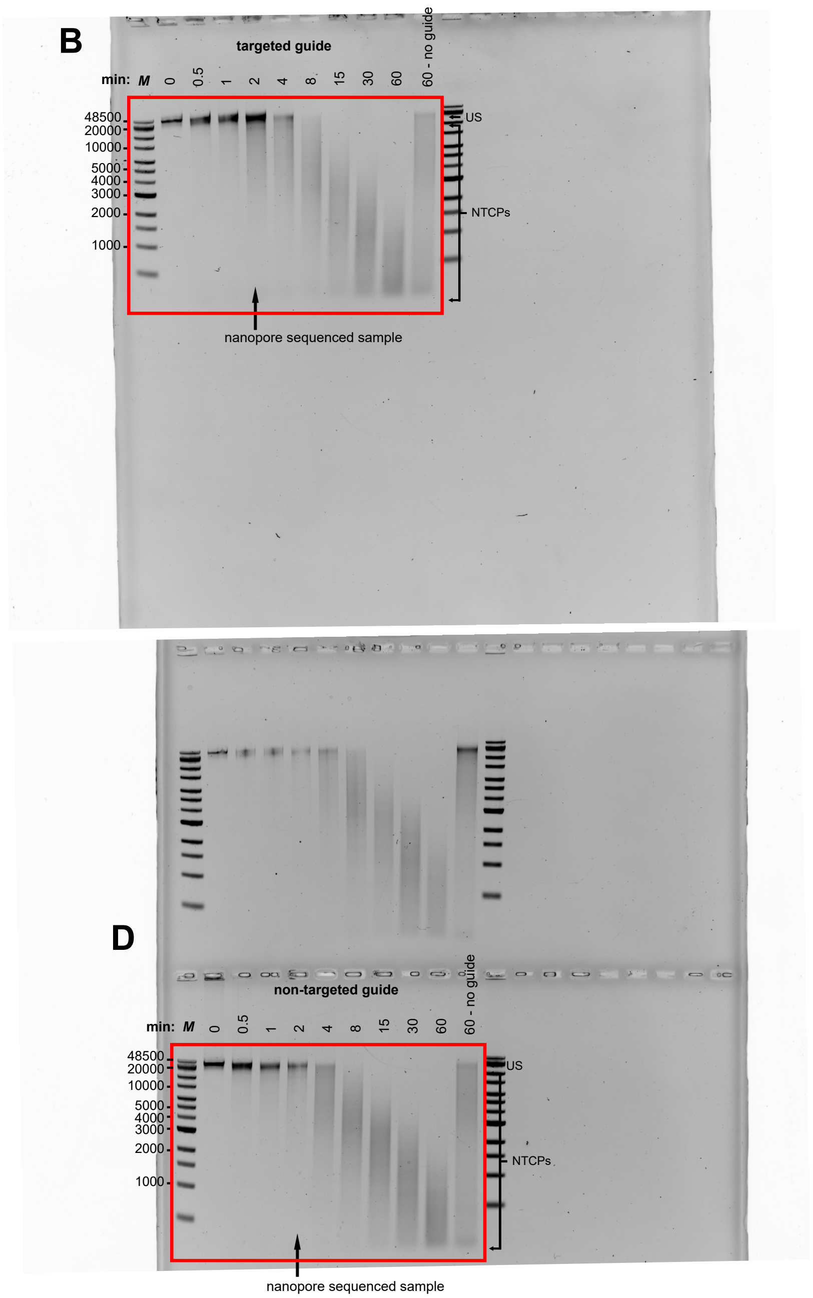


Figure S10: Uncropped agarose gels corresponding to Fig. 4 B–D. Complete gel images for $\lambda$-DNA cleavage time-courses with TevCas12a and targeted ($gRNA_{\lambda}$) or non-targeted ($gRNAA_{AAVS1}$) guides. The lane used for Oxford Nanopore sequencing is marked for reference. US = uncleaved substrate; NTCPs = non-targeted cleavage products.

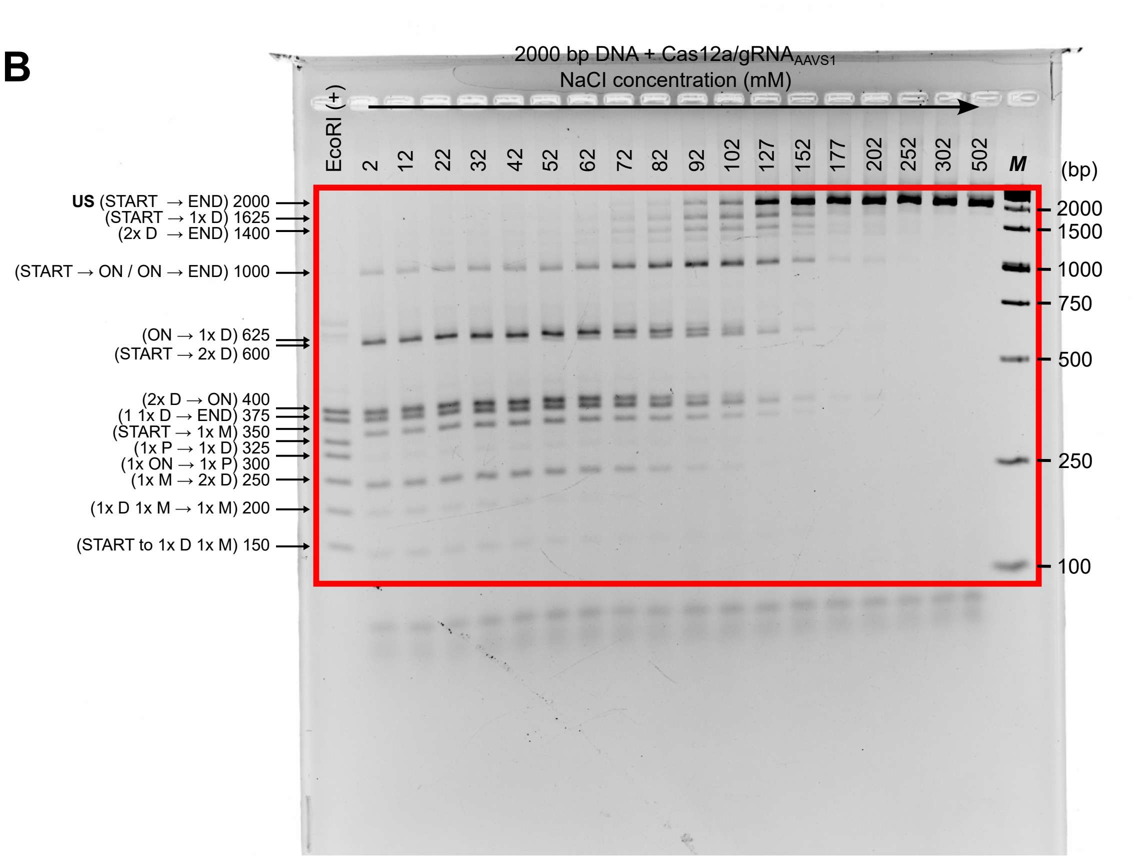


Figure S11: Uncropped agarose gel corresponding to main text **Fig. 6 B**. Full gel image of Cas12a cleavage of the 2-kb Cas-OFF substrate across increasing NaCl concentrations. Product identities and fragment sizes are as indicated in the main figure for reference. US = uncleaved substrate.

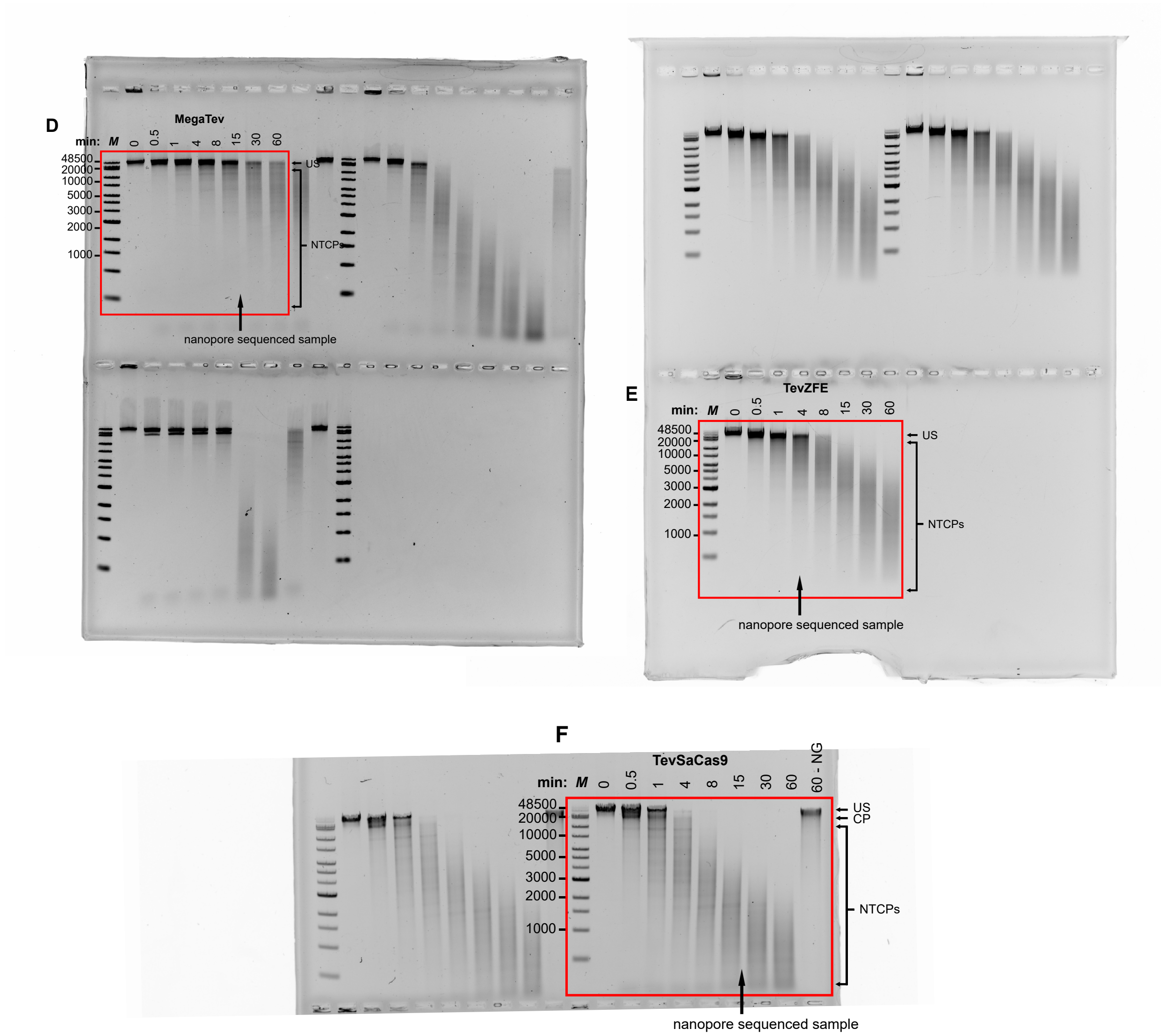


Figure S12: Uncropped agarose gels corresponding to main text **Fig. 7 D–F**. Complete time-course gels for $\lambda$-DNA cleavage by TevZFE, MegaTev, and TevSaCas9/gRNA$_\lambda$ fusions. Lanes correspond to the main-figure panels; the lane used for Oxford Nanopore sequencing is annotated for reference. US = uncleaved substrate; NTCPs = non-targeted cleavage products; CP = Cas-targeted product.

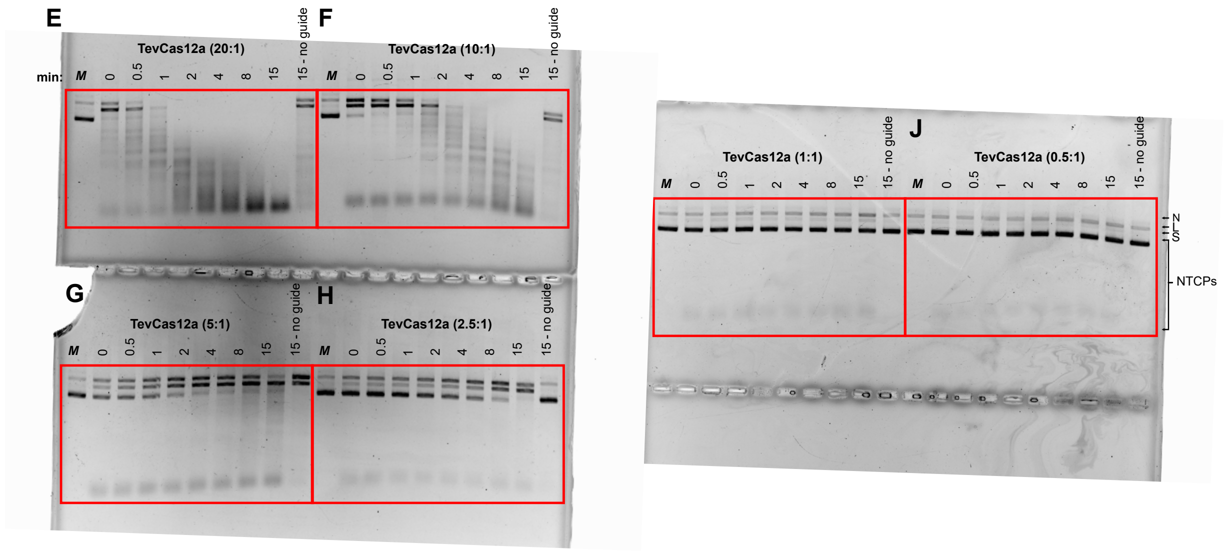


Figure S13: Uncropped agarose gels corresponding to **Supplementary Fig. S1 E–J**. Full gels showing TevCas12a cleavage of pJET-AAVS1 at varying RNP:DNA ratios (20:1 to 0.5:1). Bands for supercoiled, nicked, and linear plasmid forms are indicated for reference. NG = no guide control.

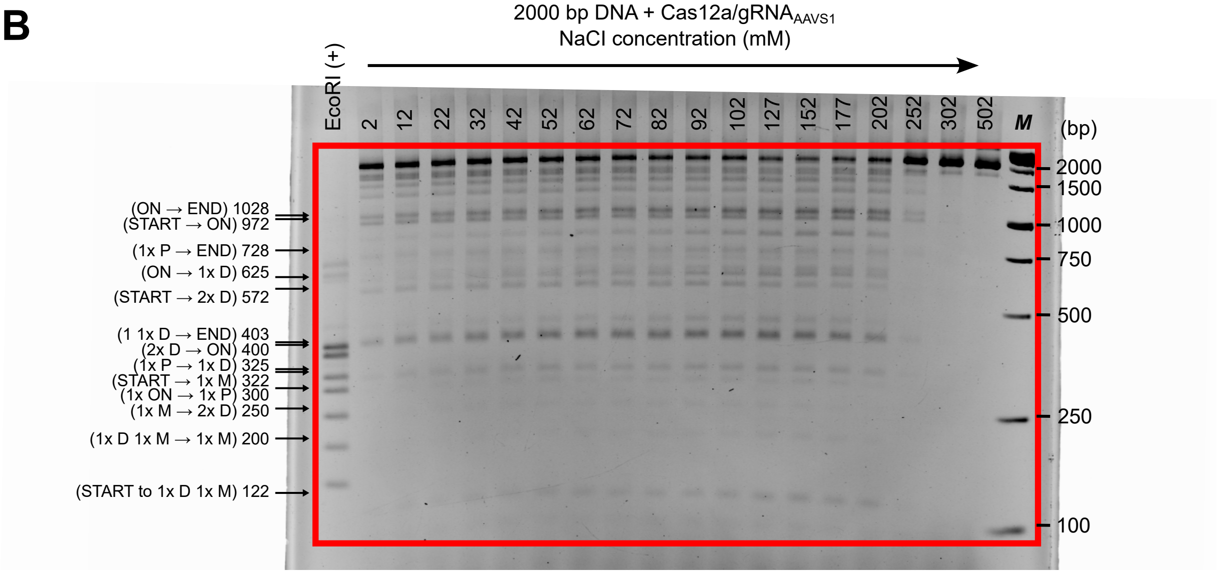


Figure S14: Uncropped agarose gel corresponding to **Supplementary Fig. 5 B**. Full gel image of SaCas9 cleavage of the 2-kb Cas-OFF substrate across increasing NaCl concentrations. Product identities and fragment sizes are as indicated in the supplementary figure for reference. US = uncleaved substrate.

### Supplementary Tables

#### Oligonucleotides and gBlocks

Table S1: Oligonucleotide and gBlock sequences and descriptions.

| Primer ID | Description | Sequence |
|---|---|---|
| OL0345 | Forward primer for amplifying AAVS1 safe harbor locus and for blunt-cloning in pJet | GCAGCTTCCTTACACTTCCC |
| OL0346 | Reverse primer for amplifying AAVS1 safe harbor locus and for blunt-cloning in pJet | CTAGGACTGAGGGTTTCAGTGC |
| OL0524 | Forward oligo for Cas12a AAVS1 gRNA IVT template | agcTAATACGACTCACTATAGTAATTTCTACTCTTGTAGATAGTCCTTGGCAAGCCCAGGAGTTTTTTT |
| OL0525 | Reverse oligo for Cas12a AAVS1 gRNA IVT template | aaaaaaaCTCCTGGGCTTGCCAAGGACTATCTACAAGAGTAGAAATTACTATAGTGAGTCGTATTAGCT |
| OL0620 | Forward primer for Cas12a for D908P substitution | CCAGAGACGCCGATCATCGGCATCCCGCGGGGTGAGCGTAACTTGATC |
| OL0621 | Reverse primer for Cas12a for D908P substitution | GCTCACCCCGCGGGATGCCGATGATCGGCGTCTC |
| OL0687 | Forward oligo for Cas12a CFTR gRNA IVT template | agcTAATACGACTCACTATAGTAATTTCTACTCTTGTAGATGAGTGATACCACAGGTGAGCTTTTTTT |
| OL0688 | Reverse oligo for Cas12a CFTR gRNA IVT template | aaaaaaaGCTCACCTGTGGTATCACTCATCTACAAGAGTAGAAATTACTATAGTGAGTCGTATTAGCT |
| DE5841 | Tev nuclease domain sequencing primer for pET11a | GTTACGGCATTTGCTGCAGG |
| DE5931 | Cas12a D908P forward sequencing primer in pET11a | AGGCACGCGCACTGCTTCCG |
| DE6018 (OL256) | SaCas9 Universal gRNA bottom strand in vitro transcription template | aaaaTCTCGGCCAACAAGTTGACGAGATAAACACGGCATTTTGCCTTGTTTTAGTAGTTCTGTTTCCAGAGGTACTAAAAC |
| DE6502 | Forward primer to amplify AAVS1 gBlocks | CGTGCGTCAGCTTTACCTGT |

Continued on next page

| Primer ID | Description | Sequence |
|---|---|---|
| DE6503 | Reverse primer to amplify AAVS1 gBlocks | GGGCTGGTGTGGCCTCTCG |
| DE6518 | Forward sequencing primer for AAVS1 inserts in pJet | CGACTCACTATAGGGAGAGCGG C |
| DE6519 | Reverse sequencing primer for AAVS1 inserts in pJet | AAGAACATCGATTTTCCATGGC AG |
| DE6524 | Forward primer for amplifying CNNNG-deletion gBlock 1 | CGTGCGTCAGTTTACCTGTG |
| DE6525 | Reverse primer for amplifying CNNNG-deletion gBlock 1 | TAGACAGGGCTGGGGTGG |
| DE6782 | Forward primer for amplifying TTTV-deletion gBlock 2 | CGTGCGTCAGTGTTACCTGTG |
| DE6783 | Reverse primer for amplifying TTTV-deletion gBlock 2 | GGACTGGGATGGCCTCTCG |
| DE6784 | Forward primer for amplifying CNNNG-TTTV-deletion gBlock 3 | CGTTCGTCAGCTGTACATGTAA G |
| DE6785 | Reverse primer for amplifying CNNNG-TTTV-deletion gBlock 3 | GGGCTGGTGTGGACTATC |
| DE7110 | Cas12a guide lambda 1 sense | agcTAATACGACTCACTATAGtaat ttctactcttgtagatTGATATGCCGCAG AAACGTTGttttttt |
| DE7111 | Cas12a guide lambda 1 anti-sense | aaaaaaaCAACGTTTCTGCGGCATA TCAatctacaagagtagaaattaCTATAGT GAGTCGTATTAgct |
| DE7112 | Cas12a guide lambda 2 sense | agcTAATACGACTCACTATAGtaatt tctactcttgtagatGCGACCTCTTCGTA CGGCCTCttttttt |
| DE7113 | Cas12a guide lambda 2 antisense | aaaaaaaGAGGCCGTACGAAGAGG TCGCatctacaagagtagaaattaCTATAG TGAGTCGTATTAgct |
| DE7114 | Cas12a guide lambda 3 sese | agcTAATACGACTCACTATAGtaatt tctactcttgtagatTGCATCCATCTGGA TTCTCCttttttt |
| DE7115 | Cas12a guide lambda 3 antisense | aaaaaaaGGAGAATCCAGATGGAT GCAatctacaagagtagaaattaCTATAGT GAGTCGTATTAgct |
| DE7182 | Unlabeled forward amplicon primer (pSP72/directionality gBlocks/CAS-OFF) | AGCAGATTGTACTGAGAGTGC |

Continued on next page

| Primer ID | Description | Sequence |
|---|---|---|
| DE7183 | 5' 6-FAM labeled forward amplicon primer (pSP72/directionality gBlocks/CAS-OFF) | /56-FAM/AGCAGATTGTACTGAGAGTGC |
| DE7184 | Unlabeled reverse amplicon primer (pSP72/directionality gBlocks/CAS-OFF) | AAGAGCGCCCAATACGC |
| DE7185 | 5' 6-FAM labeled reverse amplicon primer (pSP72/directionality gBlocks/CAS-OFF) | /56-FAM/AAGAGCGCCCAATACGC |
| DE7330 | Lambda Site 1 SaCas9 Guide Oligo for IVT | aagcTAATACGACTCACTATAGGAACCAGCAGGCGGACTTCCGGTTTTAGTACTCTGGAAACAG |
| DE7331 | Lambda Site 2 SaCas9 Guide Oligo for IVT | aagcTAATACGACTCACTATAGAATTTATAACCGACCCCAACGGTTTTAGTACTCTGGAAACAG |
| DE7332 | Lambda Site 3 SaCas9 Guide Oligo for IVT | aagcTAATACGACTCACTATAGCACACCCCAAAGCCTTCTGCTGTTTTAGTACTCTGGAAACAG |
| gBlock 1 | Sequence of modified AAVS1 with no off-target CNNNG motifs | CGTGCGTCAGCTTTACCTGTGAGATAAGGCCAGTAGCCAGACCCGTACTGGAAGGGATGTGGTGAGGAGGGGGGTGTACGTGTGGAAAACTCCATTTGTGAGAATGGTGCGTAATAGGTGTTCAACAGGTAGTGGCCGCCTCTACTCCCTTTCTCTTTCTCCATCCTTCTTTCCTTAAAGAGTCACAAGTGCTATATGGGACATATTCATCCGACAAGAGAAGGGTCCCGCTTCCATAAGGCCCTGATATGGGCTTATGGGTTTGAGTCCTTGGCAAGCCCAGGAGAGGCGATCAGGCTTACCTGTCCCCCTTACTCGTCCACCATCTCATGCAACTGGCTCTCCTGCCCCTTCCCTAAAGGGGTTACTGGATCTGCTCTTCAGAATGAGACCCGTTCACCTGCATACCCGTTCACCTGCATCCCCCTTCACCTGCATCCAAAAACGGCAACAGGCCACCTAATTGGACTGGACCACACGAGAGGCCACACCAGCCC |

Continued on next page

| **Primer ID** | **Description** | **Sequence** |
|---|---|---|
| gBlock 2 | Sequence of AAVS1 with no off-target TTTV motifs | CGTGCGTCAGTGTTACCTGTGA<br>GATAAGGCCAGTAGCCAGCCCC<br>GTCCTGGCAGGGCTGTGGTGAG<br>GAGGGGGGTGTCCGTGTGGATA<br>ACTCCCTATGTGAGAATGGTGC<br>GTCCTAGGTGTTCACCAGGTCG<br>TGGCCGCCTCTACTCCCATTCTC<br>GTTCTCCATCCTTCTGTCCTTGA<br>AGAGTCCCCAGTGCTATCTGGG<br>ACATATTCCTCCGCCCAGAGCA<br>GGGTCCCGCTTCCCTAAGGCCC<br>TGCTCTGGGCTTCTGGGTTTGA<br>GTCCTTGGCAAGCCCAGGAGAG<br>GCGCTCAGGCTTCCCTGTCCCCC<br>TTCCTCGTCCACCATCTCATGCC<br>CCTGGCTCTCCTGCCCCTTCCCT<br>ACAGGGGTTCCTGGCTCTGCTC<br>TTCAGACTGAGCCCCGTTCTCC<br>TGAATGCCCGTTCCACTGCATC<br>CCCCTTCGCCTGCATCCCACAGA<br>GGCTCCAGACCACCTACTTGGC<br>CTTGACTCCACGAGAGGCCATC<br>CCAGTCC |

Continued on next page

| **Primer ID** | **Description** | **Sequence** |
|---|---|---|
| gBlock 3 | Sequence of modified AAVS1 with no off-target TTTV or CNNNG motifs | CGTTCGTCAGCTGTACATGTAA<br>GATAAGGCCAGTAGTCAGACCT<br>GTACTGGAAGGGATGTGGTGAG<br>GAGGGGGGTGTACGTGTGGAGA<br>ACTCCAGTTGTGAGAATTGTGC<br>GTAATAGGTGTTCAACAGGTAA<br>TGGCCTTCTCTACTCCCTATCTC<br>GTTCTCCATCCTTCTGTCATTAG<br>ATAGTCATAAGTGCTATATCGG<br>ACATATTCATCCGACAAGAGAA<br>CGGTCGTGCTTACATAACGCCC<br>TGCTCTGGGCTTCTGGGTTTGA<br>GTCCTTGGCAAGCCCAGGAGAG<br>GTGATCAGGCTTACCTGTCCTC<br>CTTACTCGTCCACCATCTCATTC<br>AACTGGCTGTCCTTCCCCTTCTC<br>TAGAGGGCTTACTAGATCTGCT<br>CTTCAGAATGAGAACAGTTCAC<br>CTGCATACCCGTTCACCTTCATC<br>CCCCTTCACCTGCATACAAGAAC<br>GCCAACAGTCCACCTAATTGGA<br>CTTGACCACACGATAGTCCACA<br>CCAGCCC |
| gBlock 4 | 2000 bp Cas-OFF Fragment 1 BsaI-tailed | AGCAGATTGTACTGAGAGTGCA<br>ATACTCATACTCTTCCTTTTTCA<br>ATATTATTGAAGCATTTATCAG<br>GGTTATTGTCTCATGAGCGGAT<br>ACATATTTGAATGTACTCTGCT<br>CTGGGCTTCTGGGTTTGAGTCC<br>TTGGCAGGCCCAGGGGGAATTC<br>CACCTGACGTCTAAGAAACCAT<br>TATTATCATGACATTAACCTATA<br>AAAATAGGCGTATCACGAGGCC<br>GCCCCTGCAGCCGAATTATATT<br>ATTTTTGCCAAATAATTTTTAAC<br>AAAAGCTCTGAAGTCTTCTGAG<br>ACCAAGGCTTCTC |

Continued on next page

| Primer ID | Description | Sequence |
|---|---|---|
| gBlock 5 | 2000 bp Cas-OFF Fragment 2 BsaI-tailed | GACTGATGCTCAGTGAGTTACTACGCAGTCACTCATCCAGCATTGGGTCTCTCTTCTTCATTTAAATTCTTAGATCCCTGCTCCGGGCTTCTGGGTTTGAGTCCTTGGCAAACCCAGGAGGAATTCGTTCTAAACCATTTTTTTATTGTTGTATTATCTCTAATCTTACTACTCGATGAGTTTTCGGTATTATCTCTATTTTTAACTTGGAGAGACCAAGGCTTCTCTATCTGGTGATACATGAACAGATCCGTGCACCGTC |
| gBlock 6 | 2000 bp Cas-OFF Fragment 3 BsaI-tailed | TCCAGCATTGGGTCTCCTTGGAGCAGGTTCCATTCATTGTTTTTTTCATCATAGTGAATAAAATCAACTGCTTTAACACTTGTGCCTGAACACCATATCCATCCGGCGTAATACGACTCACTATAGGGAGAGCGGCCTCGGCTCTGGGCTTCTGGGTTTGAGTCCTTGGCAAGCCCAAGGGGAATTCCACTTCCCAAGAGGAGAAGCAGTTTGGAAAAACAAAATCAGAATAAGTTGGTCCTGAGTTCTAACTTTGGCTCTTCACCTTTCTAGTCCCCAATTTATATTGTTCCTCCGTGCGTCAGTTTTACCTGTGAGATAAGGCCAGTAGCCAGCCCCGTCCTGGCAGGGCTGTGGTGAGGAGGAGACCAAGGCTTCTC |

Continued on next page

| Primer ID | Description | Sequence |
| --- | --- | --- |
| gBlock 7 | 2000 bp Cas-OFF Fragment 4 BsaI-tailed | TCCAGCATTGGGTCTCGAGGAG GGGGGTGTCCGTGTGGAAAACT CCCTTTGTGAGAATGGTGCGTC CTAGGTGTTCACCAGGTCGTGG CCGCCTCTACTCCCTTTCTCTTT CTCCATCCTTCTTTCCTTAAAGA GTCCCCAGTGCTATCTGGGACA TATTCCTCCGCCCAGAGCAGGG TCCCGCTTCCCTAAGGCCCTGCT CTGGGCTTCTGGGTTTGAGTCC TTGGCAAGCCCAGGAGGAATTC AGGCGCTCAGGCTTCCCTGTCC CCCTTCCTCGTCCACCATCTCAT GCCCCTGGCTCTCCTGCCCCTTC CCTACAGGGGTTCCTGGCTCTG CTCTTCAGACTGAGCCCCGTTCC CCTGCATCCGAGACCAAGGCTT CTC |

Continued on next page

| Primer ID | Description | Sequence |
|---|---|---|
| gBlock 8 | 2000 bp Cas-OFF Fragment 5 BsaI-tailed | GACTGATGCTCAGTGAGTTACTACGCAGTCACTCAGGTCTCGCATCCCCGTTCCCCTGCATCCCCCTTCCCCTGCATCCCCCAGAGGCCCCAGGCCACCTACTTGGCCTGGACCCCACGAGAGGCCACCCCAGCCCTGTCTACCAGGCTGCCTTTTGGGTGGATTCTCCTCCAACCCTGCTCTGGGCTTCTAGGTTTGAGTTCTTGGCAAGCCCAGGAGGAATTCTGGAGCATTGGGGTGGGCTGGGGTTCAGAGAGGAGGGATTCCCTTCTCAGGTTACGTGGCCAAGAAGCAGGGGAGCTGGGTTTGGGTCAGGTCTGGGTGTGGGGTGACCAGCTTATGCTGTTTGCCCAGGACAGCCTAGTTTTAGCACTGAAACCCTCAGTCCTAGATCTTTCTAGAAGATCTCCTACAATATTCTCAGCTGCCATGGAAAATCGATGTTCTTCTTTTATTCTCTCAAGATTTTCAGGCTGTATATTAAAACTTATATTAAGAACCCCGCTCTGGGCTTCTGGGTTTGAGTCCTTGGCAAGCCCATGAGGAATTCGCAATCGACTCTCATGAAAACTACGAGCTAAATATTCAATATGTTCCTCTTGACCAACTTTATTCTGCATTTTTTTTGAACGAGGTTTAGAGCAAGCTTCAGGAAACTGAGACAGGAATTTTATTAAAAATTTAAATTTTGAAGAAAGTTCAGGGTTAATAGCATCCATTTTTTGCTTTGCAAGTTCCTCAGCATTCTTAACAAAAGACGTCTCTTTTGACATGTTTAAAGTTTAAACCTCCTGTGTGAAATTGTTATCCGCTCACAATTCCACACATTATACGAGCCGGAAGCATAAAGTGTAAAGCCTGGGGTGCCTAATGAGTGAGCTAACTCACATTAATTGCGTTGCGGCGTATTGGGCGCTCT |

#### Plasmid Information

Table S2: Plasmids used in this study. Plasmid and protein sequences are provided in separate FASTA files as Supplementary Data.

| Name | Backbone Vector | Description | Length | Marker |
|---|---|---|---|---|
| DE479 | pACYC Duet-1 | pTZ201D - Tev (1-201) + RyA Zinc Finger cloned NcoI/XhoI with 6x HIS tag | 4634 | Cm |
| DE708 | pACYC Duet-1 | Tev (1-169) + E2 I-OnuI with GGSGGS linker and 6x HIS tag | 5165 | Cm |
| DE4446/P0199 | pET11a | Cas12a with 3x HA, nucleoplasmin NLS, Tev protease site and 6x HIS tag | 9811 | Amp |
| DE4447/P0200 | pET11a | Tev[WT]-Cas12a with GGSGGTGGSG hinge, 3x HA, nucleoplasmin NLS, Tev protease site and 6x HIS tag | 10348 | Amp |
| DE4510/P0201 | pET11a | Tev (1-169) + SaCas9 with GGSGGTGGSG hinge, SV40 NLS, 3x HA, Tev protease site and 6x HIS tag | 9522 | Amp |
| DE4528 | pET11a | Cas12a [D908P] with 3x HA, nucleoplasmin NLS, Tev protease site and 6x HIS tag | 9811 | Amp |
| DE4530 | pET11a | TevCas12a [D908P] with GGSGGTGGSG hinge, 3x HA, nucleoplasmin NLS, Tev protease site and 6x HIS tag | 10348 | Amp |
| DE4628 | pET11a | Tev[R27A]Cas12a with GGSGGTGGSG hinge, 3x HA, nucleoplasmin NLS, Tev protease site and 6x HIS tag | 10348 | Amp |
| DE4728 | pJET | AAVS1 Safe Harbor Locus blunt-cloned in pJET | 3846 | Amp |
| DE4731 | pET11a | Tev[R27A]Cas12a[D908P] with GGSGGTGGSG hinge, 3x HA, nucleoplasmin NLS, Tev protease site and 6x HIS tag | 10348 | Amp |
| DE5032 | pET11a | Cas12a[W958A] with 3x HA, nucleoplasmin NLS, Tev protease site and 6x HIS tag | 9811 | Amp |
| DE5033 | pET11a | Tev[VKN]Cas12a[W958A] with GGSGGTGGSG hinge, 3x HA, nucleoplasmin NLS, Tev protease site and 6x HIS tag | 10348 | Amp |

#### Protein Information

Table S3: Protein constructs with their lengths and molecular weights.

| **Protein** | **Length (AA)** | **Mw (kDa)** |
|---|---|---|
| Tev (1-201) + RyA Zinc Finger | 303 | 35.041 |
| Tev (1-169) + E2 I-OnuI with GGSGGS linker and 6x HIS tag | 480 | 55.686 |
| Cas12a with 3x HA, nucleoplasmin NLS, Tev protease site and 6x HIS tag | 1390 | 160.565 |
| Tev (1-169) + Cas12a with GGSGGTGGSG hinge, 3x HA, nucleoplasmin NLS, Tev protease site and 6x HIS tag | 1569 | 180.733 |
| Tev (1-169) + SaCas9 with GGSGGTGGSG hinge, SV40 NLS, 3x HA, Tev protease site and 6x HIS tag | 1294 | 151.562 |
| Cas12a [D908P] with 3x HA, nucleoplasmin NLS, Tev protease site and 6x HIS tag | 1390 | 160.565 |
| TevCas12a [D908P] with GGSGGTGGSG hinge, 3x HA, nucleoplasmin NLS, Tev protease site and 6x HIS tag | 1569 | 180.733 |
| Tev[R27A]Cas12a with GGSGGTGGSG hinge, 3x HA, nucleoplasmin NLS, Tev protease site and 6x HIS tag | 1569 | 180.733 |
| Tev[R27A]Cas12a[D908P] with GGSGGTGGSG hinge, 3x HA, nucleoplasmin NLS, Tev protease site and 6x HIS tag | 1569 | 180.733 |

### Supplementary Code

#### Determining Terminal Position of Aligned Nanopore Reads

This Perl script parses Oxford Nanopore read alignments in SAM format output from minimap2 with -ax map-ont parameter applied to identify the terminal position of each aligned read and assess its proximity to CNNNG motifs in a reference sequence. Reads are reconstructed using CIGAR operations to determine true alignment endpoints.

```
#!/usr/bin/env perl
# Supplementary Script: motif_parser.pl
# Purpose: Parses aligned Oxford Nanopore reads (SAM format) to reconstruct aligned
    sequences,
# identify cleavage positions, and assess proximity to CNNNG motifs in a reference
    sequence.

my @reads;
my $ref = '';
my @motif_positions;

# Load reference sequence
open(my $REF, "<", "reference_sequence.fasta") or die "Could not open reference FASTA\n";
while (my $line = <$REF>) {
    chomp $line;
    $ref .= $line unless $line =~ /^>/;  # Skip FASTA header
}
close $REF;

# Identify all CNNNG motifs in reference sequence
while ($ref =~ /C...G/g) {
    push @motif_positions, $-[0];  # Store start index of motif
}

# Parse SAM alignment file
open(my $SAM, "<", "aligned_reads.sam") or die "Could not open SAM file\n";
while (my $line = <$SAM>) {
    chomp $line;
    my @fields = split /\t/, $line;

    my $read_id   = $fields[0];
    my $flag      = $fields[1];
    my $start_pos = $fields[3];
    my $cigar     = $fields[5];
    my $seq       = $fields[9];

    my @lengths = split /\D/, $cigar;
    my @ops     = split /\d+/, $cigar;
    shift @ops;  # Remove empty first element due to split

    my $pos = 0;
    my $ref_pos = $start_pos;
    my $reconstructed = '';
```

```
    for my $i (0..$#lengths) {
        my ($len, $op) = ($lengths[$i], $ops[$i]);
        if ($op eq 'M') {
            $reconstructed .= substr($seq, $pos, $len);
            $pos     += $len;
            $ref_pos += $len;
        } elsif ($op eq 'S') {
            $pos     += $len;
            $ref_pos += $len;
        } elsif ($op eq 'D') {
            $ref_pos -= $len;
        } elsif ($op eq 'I') {
            $pos     += $len;
            $ref_pos += $len;
        }
    }

    my $read_length = length($reconstructed);
    push @reads, join("\t", $read_id, $flag, $start_pos, $read_length, $reconstructed);
}
close $SAM;

# Output header
print "read_id\tstrand\tstart_pos\tread_length\tnear_CNNNG\n";

# Determine if each read start is within 5 bp of CNNNG motif
for my $read (@reads) {
    my @fields = split /\t/, $read;
    my $start = $fields[2];
    my $near_motif = 'n';
    foreach my $motif (@motif_positions) {
        if ($start > ($motif - 5) && $start < ($motif + 5)) {
            $near_motif = 'y';
            last;
        }
    }
    print join("\t", @fields[0..3], $near_motif), "\n";
}
```

Listing 1: Perl script to identify aligned read ends near CNNNG motifs. Reference sequence and SAM input filenames are generalized.

#### Extracting Cleavage Site Sequences and Frequencies

```python

import pandas as pd
from Bio import SeqIO

# Function to count positions and extract sequences
def extract_sequences(input_csv, reference_fasta, output_csv):
    # Read the input CSV into a DataFrame
    df = pd.read_csv(input_csv)

    # Filter out positions lower than 75 and not between certain ranges
    df = df[(df['position'] >= 75)]  # optionally: & (~df['position'].between(#####, #####))

    # Read the reference FASTA sequence
    reference_seq = SeqIO.read(reference_fasta, "fasta")

    # Create a dictionary to store positions and their corresponding sequences
    position_sequences = {}

    for _, row in df.iterrows():
        position = int(row['position'])
        start_pos = max(0, position - 10)  # Retain 10 nucleotides downstream
        end_pos = min(len(reference_seq), position + 11)  # Retain 10 nucleotides upstream

        # Extract the sequence and store it in the dictionary
        sequence = str(reference_seq.seq[start_pos:end_pos])
        position_sequences[position] = sequence

    # Create a DataFrame with "position," "frequency," and "sequence" columns
    position_counts = df['position'].value_counts().reset_index()
    position_counts.columns = ['position', 'frequency']
    position_counts['sequence'] = [position_sequences[position] for position in position_counts['position']]

    # Save the results to the output CSV
    position_counts.to_csv(output_csv, index=False)

if __name__ == "__main__":
    input_csv = "positions.csv"  # Input CSV of cleavage positions
    reference_fasta = "reference.fasta"  # Reference sequence in FASTA format
    output_csv = "cleavage_frequencies.csv"  # Output CSV file

    extract_sequences(input_csv, reference_fasta, output_csv)
```

Listing 2: Python script to extract cleavage site sequences and count frequencies based on reference coordinates. Input/output filenames are generalized for publication.

#### Filtering Cleavage Site Sequences for FASTA Export

This Python script filters sequences from a cleavage position dataset by frequency and reference position range, exporting each unique sequence once.

```python
import pandas as pd

# Function to filter sequences by frequency and position range and save to FASTA
def filter_and_save_to_fasta(input_csv, output_fasta, num_most_frequent=None,
    position_range=(XXXXX, XXXXX)):
    df = pd.read_csv(input_csv)
    df = df.sort_values(by='frequency', ascending=False)

    if num_most_frequent is not None:
        df = df.head(num_most_frequent)

    filtered_df = df[~df['position'].between(*position_range)]

    fasta_sequences = []
    for _, row in filtered_df.iterrows():
        position = int(row['position'])
        sequence = row['sequence']
        header = f'>Position_{position} Frequency_{row["frequency"]}\n'
        fasta_sequence = header + sequence + '\n'
        fasta_sequences.append(fasta_sequence)

    with open(output_fasta, "w") as output_handle:
        output_handle.writelines(fasta_sequences)

if __name__ == "__main__":
    input_csv = "input_frequencies.csv"
    output_fasta = "filtered_sequences.fasta"
    num_most_frequent = XXX
    position_range = (XXXXX, XXXXX)

    filter_and_save_to_fasta(input_csv, output_fasta, num_most_frequent, position_range)
```

Listing 3: Python script to filter and export cleavage site sequences to FASTA format.

#### Expanding FASTA Output Based on Observed Frequencies

This variant script writes each sequence multiple times based on observed cleavage frequency, which may be useful for downstream modeling or enrichment analyses.

```python
import pandas as pd

# Function to replicate sequences by frequency and save to FASTA
def filter_and_save_to_fasta(input_csv, output_fasta, num_most_frequent=None,
    position_range=(XXXXX, XXXXX)):
    df = pd.read_csv(input_csv)
    df = df.sort_values(by='frequency', ascending=False)

    if num_most_frequent is not None:
        df = df.head(num_most_frequent)

    filtered_df = df[~df['position'].between(*position_range)]

    fasta_sequences = []
    for _, row in filtered_df.iterrows():
        position = int(row['position'])
        sequence = row['sequence']
        frequency = int(row['frequency'])
        for i in range(1, frequency + 1):
            header = f'>Position_{position}_Frequency_{frequency}_{i}\n'
            fasta_sequence = header + sequence + '\n'
            fasta_sequences.append(fasta_sequence)

    with open(output_fasta, "w") as output_handle:
        output_handle.writelines(fasta_sequences)

if __name__ == "__main__":
    input_csv = "input_frequencies.csv"
    output_fasta = "replicated_sequences.fasta"
    num_most_frequent = XXX
    position_range = (XXXXX, XXXXX)

    filter_and_save_to_fasta(input_csv, output_fasta, num_most_frequent, position_range)
```

Listing 4: Python script to export sequences to FASTA format according to cleavage frequency.

#### Plotting Cleavage Frequencies on a Logarithmic Scale

This Python script reads a CSV file with cleavage site positions and frequencies, highlights sequences matching known motifs or within a specific region, and produces a log-scale scatter plot saved as a PDF.

```

import pandas as pd
import matplotlib.pyplot as plt
import numpy as np
import re

# Function to check for motifs and generate a log-scale graph with colored points and borders
def generate_frequency_graph(input_csv, output_pdf):
    # Read the CSV into a DataFrame
    df = pd.read_csv(input_csv)

    # Sort the DataFrame by 'frequency' column in descending order and select the top 250 rows
    df_sorted = df.sort_values(by='frequency', ascending=False).head(250)

    # Define regular expressions for the motifs
    motif1_pattern = r'C\w\w\wG'  # "CNNNG" motif
    motif2_pattern = r'Z\w\w\wZ'  # Optional/alternative motif placeholder

    # Create a list to store colors based on motif presence or range
    point_colors = []
    point_borders = []

    # Iterate through the rows and determine colors, borders based on motifs and range
    for _, row in df_sorted.iterrows():
        position = row['position']
        sequence = row['sequence']
        if position >= XXXXX and position <= XXXXX:
            point_colors.append('#0072b2')  # Color for the range
            point_borders.append('#00588a')  # Darker border color
        elif re.search(motif1_pattern, sequence) or re.search(motif2_pattern, sequence):
            point_colors.append('#e69f00')  # Color for motifs
            point_borders.append('#b87d00')  # Darker border color
        else:
            point_colors.append('#76c2ed')  # Default color
            point_borders.append('#5394bf')  # Darker border color

    # Create the plot with a logarithmic y-axis scale and colored borders
    plt.figure(figsize=(10, 6))
    plt.scatter(df_sorted['position'], df_sorted['frequency'], color=point_colors,
    edgecolor=point_borders)

    # Customize the plot
    plt.yscale('log')  # Set y-axis to logarithmic scale
    plt.title('Log-Scale Frequency vs. Position in Lambda', fontname='Arial',
    fontsize=14)
    plt.xlabel('Position in Lambda', fontname='Arial', fontsize=12)
```

```
    plt.ylabel('Cleavage Product Frequency', fontname='Arial', fontsize=12)

    # Save the graph as a PDF file
    plt.savefig(output_pdf, format='pdf', bbox_inches='tight')
    plt.show()

if __name__ == "__main__":
    input_csv = "cleavage_frequencies.csv"  # Replace with your input CSV file name
    output_pdf = "log_scale_frequency_graph.pdf"  # Replace with your desired output PDF
    file name

    generate_frequency_graph(input_csv, output_pdf)
```

Listing 5: Python script for plotting cleavage frequencies by position on a logarithmic scale.

#### Identifying TTTV PAM Sites in Reference Sequences

This Python script searches a reference sequence (e.g., Lambda phage) for TTTV PAM sites (where V = A/C/G) and writes the start and end positions of each match to a CSV file. To capture PAMs on the reverse strand, the script should be rerun on the reverse complement of the reference sequence, or with a reverse complement PAM input.

```python
from Bio import SeqIO
import re
import csv

# Define the PAM pattern (TTTV)
pam_pattern = re.compile(r'TTT[AGC]')

# Define the reference sequence file
reference_file = "reference_sequence.fasta"  # Generalized for publication

# Read the reference sequence
reference_sequence = SeqIO.read(reference_file, "fasta").seq

# Find PAM positions in the sequence
pam_positions = []
for match in pam_pattern.finditer(str(reference_sequence)):
    start_position = match.start() + 1  # Convert to 1-based indexing
    end_position = match.end()
    pam_positions.append((start_position, end_position))

# Define the CSV file name
csv_file = "TTTV_PAM_positions.csv"

# Write PAM positions to a CSV file
with open(csv_file, "w", newline="") as csvfile:
    csv_writer = csv.writer(csvfile)

    # Write header row
    csv_writer.writerow(["PAM", "Start Position", "End Position"])

    # Write PAM positions
    for i, (start, end) in enumerate(pam_positions, 1):
        csv_writer.writerow([f"PAM {i}", start, end])

print(f"PAM positions saved to {csv_file}")
```

Listing 6: Python script to identify TTTV PAM motifs in a reference sequence. Run separately on the reverse complement to capture both strands.

#### Calculating Distance to Nearest PAM Sites

This Python script determines the closest upstream and downstream PAM motifs (e.g., BAAA or TTTV) relative to mapped cleavage positions. It excludes reads within a defined range (if specified) and outputs separate CSV files for upstream and downstream distances.

```python

import pandas as pd

# Load PAM positions and cleavage data into DataFrames
baaa_df = pd.read_csv('BAAA_pam_positions.csv')             # Replace with your BAAA motif CSV
tttv_df = pd.read_csv('TTTV_pam_positions.csv')             # Replace with your TTTV motif CSV
barcode_df = pd.read_csv('cleavage_positions.csv')          # Replace with your cleavage data CSV

# Initialize empty lists to store distance calculations
end_distances_data = []
start_distances_data = []

# Iterate through each cleavage position
for position in barcode_df['position']:
    # Skip positions in exclusion range
    if XXXXX <= position <= XXXXX:
        continue

    # Find the nearest upstream BAAA end position
    nearest_end_positions = baaa_df['End Position'][baaa_df['End Position'] < position]
    if not nearest_end_positions.empty:
        nearest_end_position = nearest_end_positions.max()
        end_distance = position - nearest_end_position
        end_distances_data.append([position, nearest_end_position, end_distance])

    # Find the nearest downstream TTTV start position
    nearest_start_positions = tttv_df['Start Position'][tttv_df['Start Position'] > position]
    if not nearest_start_positions.empty:
        nearest_start_position = nearest_start_positions.min()
        start_distance = nearest_start_position - position
        start_distances_data.append([position, nearest_start_position, start_distance])

# Convert results to DataFrames and export to CSV
end_distances_df = pd.DataFrame(end_distances_data, columns=['position', 'nearest_end_position', 'distance'])
start_distances_df = pd.DataFrame(start_distances_data, columns=['position', 'nearest_start_position', 'distance'])

end_distances_df.to_csv('distances_to_end.csv', index=False)
start_distances_df.to_csv('distances_to_start.csv', index=False)
```

Listing 7: Python script for calculating distances from cleavage positions to nearest PAM motifs upstream (BAAA) and downstream (TTTV).

#### R Script for Plotting Distance vs. Frequency

This R script plots cleavage frequencies as a function of distance from PAMs, using smoothing and fixed axis scaling.

```
# Load required libraries
library(ggplot2)
library(dplyr)

# Replace with the actual path to your CSV file
csv_file_path <- "your_data.csv"

# Read your CSV file into a data frame
data <- read.csv(csv_file_path)

# Define specific colors for each sample value
sample_colors <- c("black", "black", "black", "black", "black")

# Create a ggplot and add layers with specific colors based on the "sample" column
ggplot(data, aes(x = distances, y = frequencies, color = as.character(sample))) +
  geom_point() +
  geom_smooth(method = "loess", se = TRUE, color = "black") +
  labs(x = "Distances", y = "Frequencies", color = "Sample") +
  scale_color_manual(values = sample_colors) +
  scale_y_continuous(limits = c(0.005, 0.0425)) +
  theme_minimal()
```

Listing 8: R script to plot frequencies as a function of distance from PAM motifs.

#### CNNNG Motif Identification and Annotation

This script identifies all CNNNG motifs in a reference sequence, extracts their positions, central triplets, calculates reverse complements, and outputs the results as a CSV.

```python

from Bio import SeqIO
import re
import csv

# Define the reference DNA sequence file and output file
input_file = "reference_sequence.fasta"
output_file = "cnnng_motif_positions.csv"

# Define the CNNNG motif pattern
motif_pattern = "C[ATGC]{3}G"

# Function to find motif positions and central triplets
def find_motif_positions(sequence, motif_pattern):
    positions = []
    central_triplets = []
    matches = re.finditer(f'(?=({motif_pattern}))', sequence)  # Overlapping matches
    for match in matches:
        start_pos = match.start(1) + 1  # Convert to 1-based
        end_pos = match.end(1)
        central_triplet = match.group(1)[1:4]
        positions.append((start_pos, end_pos))
        central_triplets.append(central_triplet)
    return positions, central_triplets

# Reverse complement function
def reverse_complement(sequence):
    complement = {'A': 'T', 'T': 'A', 'G': 'C', 'C': 'G'}
    return ''.join(complement[base] for base in reversed(sequence))

# Read sequence
with open(input_file, "r") as handle:
    sequence = str(next(SeqIO.parse(handle, "fasta")).seq)

# Find motif matches
positions, central_triplets = find_motif_positions(sequence, motif_pattern)
reverse_complement_triplets = [reverse_complement(t) for t in central_triplets]

# Save to CSV
with open(output_file, "w", newline="") as csvfile:
    writer = csv.writer(csvfile)
    writer.writerow(["Motif", "Start Position", "End Position", "Central Triplet", "Reverse Complement Triplet"])
    for i, (start, end) in enumerate(positions):
        writer.writerow([f"Motif_{i+1}", start, end, central_triplets[i], reverse_complement_triplets[i]])

print(f"Motif data saved to {output_file}")
```

Listing 9: Python script for identifying CNNNG motifs and their reverse complements from a reference sequence.

#### Motif Frequency Normalization by Triplet Occurrence for Heatmapping

This script normalizes observed motif cleavage frequencies based on the total occurrence of each central triplet within a reference genome. It must be run separately for both forward and reverse complement motif datasets.

```
import csv
import os
from Bio import SeqIO

# Input directory containing position-frequency CSV files
input_directory = "input_directory_of_position_frequency_csvs"

# Other input files
motif_positions_file = "motif_positions.csv"         # e.g.,
    forward_triplets_in_range.csv or reverse_triplets_in_range.csv
reference_sequence_file = "reference_sequence.fasta" # e.g., Lambda_NEB.fasta
triplet_counts_file = "triplet_counts.csv"           # e.g., forward_triplet_counts.csv
    or reverse_triplet_counts.csv

# Output directory (can be the same or different)
output_directory = "output_directory_for_counts"

# Ensure output directory exists
os.makedirs(output_directory, exist_ok=True)

# Dictionary to store triplet counts
triplet_counts = {}

# Read triplet counts from triplet_counts.csv
with open(triplet_counts_file, "r") as triplet_counts_data:
    csvreader = csv.DictReader(triplet_counts_data)
    for row in csvreader:
        triplet = row["Triplet"]
        count = int(row["Count"])
        triplet_counts[triplet] = count

# Read reference sequence from FASTA
with open(reference_sequence_file, "r") as handle:
    for record in SeqIO.parse(handle, "fasta"):
        reference_sequence = str(record.seq)

# Read motif positions and central triplets from motif_positions.csv
motif_data = []
with open(motif_positions_file, "r") as motif_file:
    csvreader = csv.DictReader(motif_file)
    for motif in csvreader:
        motif_data.append({
            "start_pos": int(motif["Start"]),
            "end_pos": int(motif["End"]),
            "central_triplet": motif["Triplet"]
        })

```

```python
# Process each CSV file in the input directory
for filename in os.listdir(input_directory):
    if filename.endswith(".csv"):
        file_path = os.path.join(input_directory, filename)
        output_file = os.path.join(output_directory, f"{os.path.splitext(filename)[0]}_counts.csv")

        # Dictionary to store frequencies for each central triplet in this file
        central_triplet_frequencies = {}

        with open(file_path, "r") as positions_data:
            csvreader = csv.DictReader(positions_data)
            for row in csvreader:
                try:
                    position = int(float(row["position"]))  # Convert to float and then to int
                except ValueError:
                    continue
                frequency = int(row["frequency"])

                for motif in motif_data:
                    if motif["start_pos"] <= position <= motif["end_pos"]:
                        central_triplet = motif["central_triplet"]
                        central_triplet_frequencies.setdefault(central_triplet, 0)
                        central_triplet_frequencies[central_triplet] += frequency

        # Write results for this file
        with open(output_file, "w", newline="") as output_csv:
            csvwriter = csv.writer(output_csv)
            csvwriter.writerow(["Central Triplet", "Total Frequency", "Normalized Frequency"])
            for triplet, frequency in central_triplet_frequencies.items():
                count = triplet_counts.get(triplet, 0)
                normalized_frequency = frequency / count if count > 0 else 0
                csvwriter.writerow([triplet, frequency, normalized_frequency])

        print(f"Results saved to {output_file}")

# NOTE: This script must be run separately for both forward and reverse complement motif windows.
```

Listing 10: Python script to normalize motif cleavage frequency by reference triplet abundance.
